# Computationally efficient whole genome regression for quantitative and binary traits

**DOI:** 10.1101/2020.06.19.162354

**Authors:** Joelle Mbatchou, Leland Barnard, Joshua Backman, Anthony Marcketta, Jack A. Kosmicki, Andrey Ziyatdinov, Christian Benner, Colm O’Dushlaine, Mathew Barber, Boris Boutkov, Lukas Habegger, Manuel Ferreira, Aris Baras, Jeffrey Reid, Gonçalo Abecasis, Evan Maxwell, Jonathan Marchini

**Author notes:** **Corresponding author** Jonathan Marchini.

## Abstract

Genome-wide association analysis of cohorts with thousands of phenotypes is computationally expensive, particularly when accounting for sample relatedness or population structure. Here we present a novel machine learning method called REGENIE for fitting a whole genome regression model that is orders of magnitude faster than alternatives, while maintaining statistical efficiency. The method naturally accommodates parallel analysis of multiple phenotypes, and only requires local segments of the genotype matrix to be loaded in memory, in contrast to existing alternatives which must load genomewide matrices into memory. This results in substantial savings in compute time and memory usage. The method is applicable to both quantitative and binary phenotypes, including rare variant analysis of binary traits with unbalanced case-control ratios where we introduce a fast, approximate Firth logistic regression test. The method is ideally suited to take advantage of distributed computing frameworks. We demonstrate the accuracy and computational benefits of this approach compared to several existing methods using quantitative and binary traits from the UK Biobank dataset with up to 407,746 individuals.

## Introduction

Since the first large genome-wide association studies^1^ were carried out in 2007, there has been a steady increase in sample sizes, now reaching 100,000s of individuals, enabled by a parallel stream of methods with ever increasing computational efficiency. Initial methods used simple linear regression and logistic regression using programs such as SNPTEST^1^ and PLINK^2^ (see **URLs**), but these have largely been replaced by the use of linear mixed models (LMMs) and closely related whole-genome regression (WGR) models. These approaches have been shown to account for population structure and relatedness, and offer advantages in power by conditioning on associated markers from across the whole genome^3–7^.

Initial methods were focused on quantitative traits^3^ for studies with a few thousand samples and assumed a Gaussian distribution on SNP effect sizes. These approaches were extended to datasets including 10,000s of individuals by computational strategies that avoided repeated matrix inversions when testing each SNP^7,8^. Building on work from the plant and animal breeding literature^9,10^, even more efficient whole genome regression approaches were developed that allowed for more flexible (non-Gaussian) prior distributions of SNP effect sizes^11,12^. The BOLT-LMM method is one commonly used implementation of this approach^13,14^. In addition, the fastGWA LMM approach was recently introduced which reduces the computational time by using a sparse representation of the genetic correlations present in the sample^15^. Extensions that allow for general gene-environment effects to be inferred have also been developed^16^. For simple linear regression of quantitative traits, the BGENIE method introduced the idea of simultaneous analysis of multiple quantitative traits, which required only a single pass through the genetic data and provided substantial speed-ups over PLINK^17^.

BOLT-LMM and fastGWA have also been applied to binary (case-control) traits, when the case-control ratio is reasonably balanced and relatively common variants are tested for association. However, these approaches break down when applied to unbalanced case-control studies tested and rarer variants, such as those produced through exome sequencing studies. The SAIGE method implements a logistic mixed model approach that explicitly models the binary nature of the trait^18^. When testing variants, SAIGE uses a saddle pointapproximation (SPA) to the test statistic null distribution, and this is quite effective at controlling Type I error.

The BOLT-LMM, fastGWA and SAIGE methods all proceed in two main steps that are applied one trait at a time. In Step 1, a model is fit to a set of SNPs from across the whole genome, such as all the SNPs on a genotyping array. The resulting model fit is then used to create either a prediction of individual trait values based on the genetic data (in BOLT-LMM and SAIGE) or an estimate of the trait variance-covariance matrix (in fastGWA). In Step 2, a larger set of imputed or sequenced variants on the same set of samples are tested for association, conditional upon the predictions or variance-covariance matrix in Step 1. This is usually carried out using the so-called LOCO (leave one chromosome out) scheme, where each imputed SNP on a chromosome is tested conditional on the Step 1 predictions ignoring that chromosome. This approach avoids proximal contamination, which can reduce association test power^8,19^.

In this paper, we propose a new machine learning method within this 2-step paradigm, called REGENIE, that is substantially faster than existing approaches. **Supplementary Figure S1** provides an overview of our REGENIE method. In Step 1, array SNPs are partitioned into consecutive blocks of *B* SNPs and from each block a small set of *J* ridge regression predictions are generated. Within each block, the ridge regression predictors each use a slightly different set of shrinkage parameters. The idea behind using a range of shrinkage values is to capture the unknown number and size of truly associated genetic markers within each window. This approach is equivalent to placing a Gaussian prior on the effect sizes of the SNPs in the block and finding the maximum a posteriori (MAP) estimate of the effect sizes and the resulting prediction. One can think of these predictions as local polygenic scores that account for local linkage disequilibrium (LD) within blocks. Combining the predictions together from across the genome results in a large reduction in the size of the genetic dataset. In this paper, we use B = 1000 and J = 5 and this reduces a set of *M* = 500, 000 SNPs to *M* = 2, 500 predictors. The method then uses a second ridge regression (linear or logistic depending on the phenotype) within a cross-validation (CV) scheme (either K-fold CV or leave-one-out CV) to combine the M predictors into a single predictor, which is then decomposed into 23 chromosome predictions for the LOCO approach. These are then used as a covariate in Step 2 when each imputed SNP is tested. This approach completely decouples Step 1 and Step 2, so that the Step 1 predictions can be re-used when running Step 2 on distinct sets of markers (imputed and exome markers for example) or even when distinct statistical tests at Step 2 are needed.

This approach exhibits a number of desirable properties. First, many of the calculations in Steps 1 and 2 can be carried out for multiple traits in parallel. This leads to substantial gains in speed since the files containing the SNPs in Step 1 and SNPs in Step 2 are read only once, rather than repeatedly for each trait. In practice, we find that for Step 1 REGENIE can be over 150x faster than BOLT-LMM and 300x faster than SAIGE when analyzing 50 UK Biobank quantitative and binary traits with up to 407,746 samples (**Tables 1–2**). For Step 2, the differences between methods are less extreme and depend upon the type of trait, test statistic and implementations of file format reading and parallelization schemes, but we show that REGENIE is still substantially faster than other approaches. In the **Supplementary Methods** and **Supplementary Table S1** we provide an analysis of the computational complexity of REGENIE.

**Table 1:** Comparison of runtimes of REGENIE, fastGWA, and BOLT-LMM when analyzing 50 quantitative traits with UK Biobank data. For REGENIE and BOLT LMM, 469,336 LD-pruned SNPs were used as model SNPs when fitting the null model (step 1). For fastGWA, these SNPs were used to compute the sparse GRM with the default relatedness threshold of 0.05 (step 0*). Tests were performed on 11.4M imputed SNPs (step 2). For step 1, REGENIE was run in multi-trait (MT) mode analyzing all traits together at once using 5-fold cross-validation (REGENIE-MT) as well as using leave-one out cross validation (REGENIE-LOOCV). All runs were done on the same computing environment (16 virtual CPU cores of a 2.1GHz AMD EPYC 7571 processor, 64GB of memory and 600GB solid-state disk) except for the GRM calculation required for fastGWA, where we used 250 partitions in a computing environment with 4 virtual CPU cores and 8GB of memory. The BGEN file input needed for step 2 was split by chromosome so fastGWA had to be ran separately for each chromosome being tested. The sample sizes for the 50 traits range from 332,739 to 407,662 individuals (see **Supplementary Table S2**).

| Method | Step | Benchmark |  |  |
| --- | --- | --- | --- | --- |
|  |  | CPU Time (hours) | Elapsed time (hours) | Memory usage (GB) |
| REGENIE MT | 1 | 111 | 12 | 12.9 |
| REGENIE MT-LOOCV | 1 | 192 | 19 | 23.6 |
| REGENIE MT | 2 | 1,108 | 75 | 6.0 |
| fastGWA | 0* | 454 | 201 | — |
| fastGWA | 1-2 | 3,546 | 243 | 2.0 |
| BOLT-LMM | 1 | 39,806 | 1,815 | 50 |
| BOLT-LMM | 2 |  | 863 |  |

**Table 2:** Comparison of runtimes of REGENIE-FIRTH, REGENIE-SPA, and SAIGE when analyzing 50 binary traits with UK Biobank data. 469,336 LD-pruned SNPs were used as model SNPs when fitting the null model (step 1). Tests were performed on 11.8M imputed SNPs (step 2). For step 1, REGENIE was run in multi-trait (MT) mode analyzing all traits together at once using 5-fold cross-validation (REGENIE-MT) as well as using leave-one out cross validation in step 1 (REGENIE-LOOCV). For step 2, REGENIE was run using Firth correction (REGENIE MT-FIRTH) and using saddle pointapproximation (REGENIE MT-SPA). Step 1 of SAIGE did not finish for 2/50 traits as it exceeded the 4-week limit, so SAIGE step 2 reported timings are projections based on timings of the completed runs. All runs were done on the same computing environment (16 virtual CPU cores of a 2.1GHz AMD EPYC 7571 processor, 64GB of memory and 600GB solid-state disk). The sample sizes for the 50 traits range from 381,591 to 407,746 individuals (see **Supplementary Table** S6)

| Method | Step | Benchmark |  |  |
| --- | --- | --- | --- | --- |
|  |  | CPU Time (hours) | Elapsed time (hours) | Memory usage (GB) |
| REGENIE MT | 1 | 1,590 | 117 | 11.8 |
| REGENIE MT-LOOCV | 1 | 777 | 108 | 19.5 |
| REGENIE MT-FIRTH | 2 | 45,427 | 3,240 | 7.7 |
| REGENIE MT-SPA | 2 | 31,216 | 2,002 | 9.1 |
| SAIGE* | 1 | 275,070 | 21,428 | 49 |
| SAIGE* | 2 | 94,347 | 68,437 | 2.1 |

Second, in Step 1 of REGENIE, only B SNPs need to be stored in memory at once, which leads to a low memory footprint. Third, the method is applicable to both quantitative and binary traits, and for binary traits we have implemented a new, fast Firth logistic regression test, as well as a saddle pointapproximation (SPA) test. Finally, our algorithm is ideally suited to implementation on distributed computing frameworks such as Apache Spark, where both the data set and application of the method and computation can be parallelized across a large number of machines. The main implementation of REGENIE is a standalone C++ program, but these methods have also been implemented for quantitative traits in the Glow project, which is based on Apache Spark (see **URLs**). All of the main experiments and results in the paper were obtained using the C++ program.

## Results

### Quantitative Traits

**Figure 1** shows the results of applying REGENIE, BOLT-LMM and fast-GWA to 3 quantitative phenotypes measured on white British UK Biobank participants (LDL, *N* = 389, 189; Body mass index, *N* = 407, 609; and Bilirubin, *N* = 388, 303) where Step 2 testing was performed on 9.8 million imputed SNPs (see **Supplementary Methods**). The Manhattan plots for all three phenotypes show good agreement between the methods (see also **Supplementary Figure S2**) with both REGENIE and BOLT-LMM showing increased power gains relative to fastGWA at known peaks of association.

**Figure 1:**
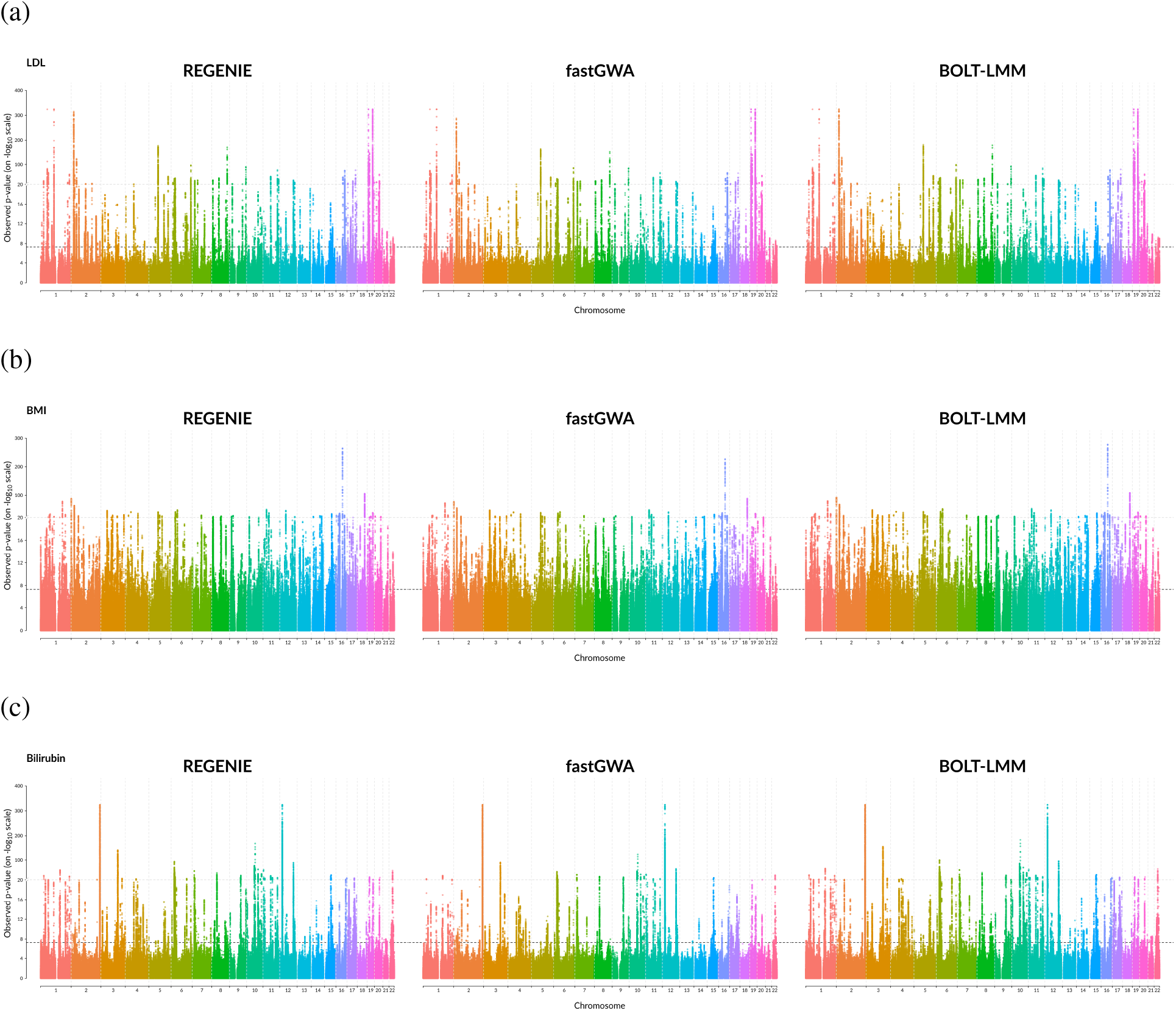
Comparison of methods on quantitative traits from UK Biobank. Results from REGENIE, fastGWA and BOLT-LMM on white British samples for (a) LDL (*N* = 389,189), (b) BMI (*N* = 407,609) and (c) Bilirubin (*N* = 388,303). Tests were performed on 9.8M imputed SNPs with minor allele frequency above 1%. The lower dashed horizontal line represents the genomewide significance level of 5 × 10^−8^ and the upper dashed horizontal line represents the breakpoint for the different scaling of the y-axis. The dashed vertical lines separate the 22 chromosomes.

To demonstrate the advantages of analyzing multiple traits in parallel using REGENIE, we compared it to BOLT-LMM and fastGWA on a set of 50 quantitative traits from the UK Biobank, each with a distinct missing data pattern (see **Supplementary Table S2**). While REGENIE can analyze all traits at once within a single run of the software, the BOLT-LMM and fastGWA software must be run once for each of the 50 traits. Across all 50 traits, we find that REGENIE and BOLT-LMM p-values are in very close agreement (see **Supplementary Figure S3**), but often the fastGWA p-values look noticeably deflated compared to REGENIE and BOLT-LMM. The computation time and memory usage of the 3 methods is given in **Table 1**. The table shows that in this 50 traits scenario REGENIE is 151x faster than BOLT-LMM in elapsed time for Step 1 and 11.5x faster for Step 2, and this translates into >30x overall speed-up in terms of elapsed time. Similar to BOLT-LMM, step 2 of REGENIE has been optimized for input genotype data in BGEN v1.2 format which helped significantly reduce the runtime. In addition, REGENIE has a maximum memory usage of 12.9 GB, which is mostly due to REGENIE only reading a small portion of the genotype data at a time, whereas BOLT-LMM required 50GB. In order to keep memory usage low when analyzing the 50 traits, within-block predictions are stored on disk and read separately for each trait working across blocks. The added I/O operations incur a small cost on the overall runtime but significantly decrease the amount of memory needed by REGENIE (see **Supplementary Table S3**). When running analyses on cloud-based services such as Amazon Web Services (AWS), these time and memory reductions both contribute to large reductions in cost, as cheaper AWS instance types can be used and for less time. In the same 50 traits scenario, we find that REGENIE is 2.8x faster than fastGWA, but fastGWA is very memory efficient and uses only a maximum of 2GB.

### Binary Traits

In addition to analyzing quantitative traits, REGENIE was also designed for the analysis of binary traits, and it is in this setting that some of the largest benefits of using REGENIE occur. Analysis of an unbalanced binary trait can lead to elevated Type 1 error if the imbalance is not accounted for in some way^18^. REGENIE includes implementations of both Firth and SPA corrections to also be able to handle this scenario (see **Methods**). **Figure 2** (see also **Supplementary Figure S4**) show the results of applying REGENIE, BOLT-LMM and SAIGE to 4 binary phenotypes measured on white British UK Biobank participants (coronary artery disease [CAD], *N* = 352, 063; glaucoma, *N* = 406, 927; colorectal cancer, *N* = 407, 746; and thyroid cancer, *N* = 407, 746) where Step 2 testing was performed on 11.6 million imputed SNPs (see **Supplementary Methods**). All four approaches show very good agreement for the most balanced trait (CAD; case-control ratio=1:11), but as the fraction of cases decreases BOLT-LMM tends to give inflated test statistics, as expected^18^. However both REGENIE with Firth and SPA corrections, as well as SAIGE, which uses SPA correction, are all robust to this inflation and show similar agreement for the associations detected.

**Figure 2:**
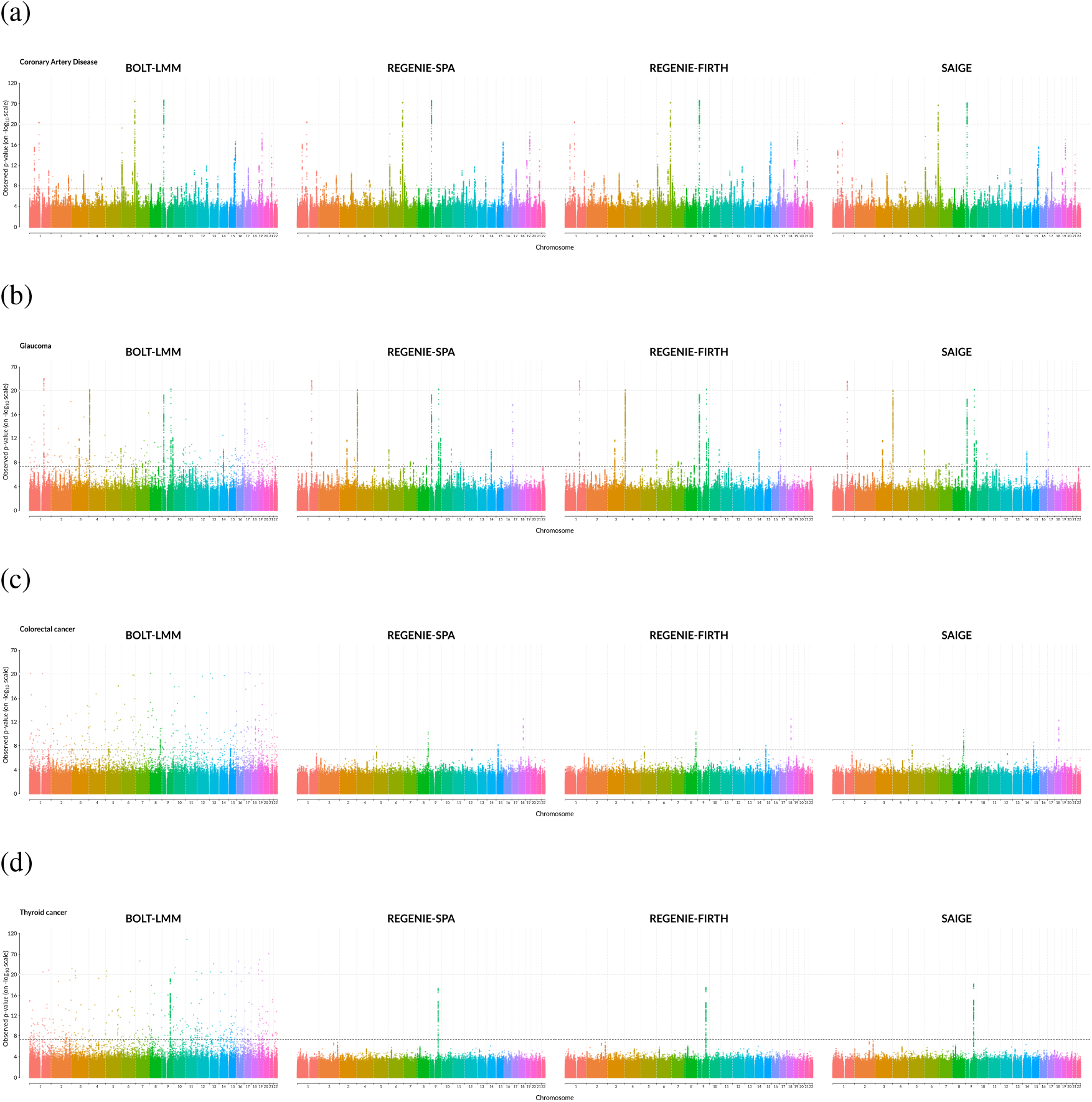
Comparison of methods on binary traits from UK Biobank. Results from REGENIE using Firth and SPA correction, BOLT-LMM and SAIGE on white British samples are shown for (a) coronary artery disease (case-control ratio=1:11, *N* = 352,063), (b) glaucoma (case-control ratio=1:52, *N* = 406, 927), (c) colorectal cancer (case-control ratio=1:97, *N* = 407, 746), and (d) thyroid cancer (case-control ratio=1:660, *N* = 407, 746). Tests were performed on 11 million imputed SNPs. The lower dashed horizontal line represents the genome-wide significance level of 5 × 10^−8^ and the upper dashed horizontal line represents the breakpoint for the different scaling of the y-axis. The dashed vertical lines separate the 22 chromosomes.

The SPA and Firth corrections are fundamentally different. The SPA approach calculates a standard score test statistic and approximates the null distribution whereas the Firth correction uses a penalized likelihood approach to estimate SNP effect size parameters in an asymptotic likelihood ratio test. While both provide good control of Type 1 error for rare binary traits, we found that SPA approach implemented in SAIGE can result in very inflated effect size estimates (see **Supplementary Figure S5**). However, the Firth correction used in REGENIE not only provides good calibration of the test, but also gives effect size estimates and standard errors which are robust to inflation due to low minor allele counts (see **Supplementary Table S4**). The fast Firth correction that we developed not only leads to similar effect sizes and p-values compared to the exact Firth correction (see **Supplementary Figures S7-S8**), but is ~ 60 times faster (see **Supplementary Table S5**).

To assess the computational resources needed with a larger number of traits being analyzed, we again ran REGENIE using Firth/SPA correction and SAIGE on a set of 50 binary traits from the UK Biobank with a range of different case-control ratios and distinct missing data patterns (see **Supplementary Table S6**). The computation time and memory usage details are given in **Table 2**.

For Step 1, we find that REGENIE (using the leave-one-out CV scheme) was about 350 times faster (777 vs 275,070 CPU hours) and required only 40% of the memory used by SAIGE (19.5GB versus 49GB). In Step 2, REGENIE Firth and SPA were 2x and 3x faster than SAIGE in CPU time, respectively, but were 21x and 34x faster than SAIGE in elapsed time, respectively, which suggests that REGENIE makes better use of parallelization in this step. Overall, in this 50 trait setting, REGENIE Firth was 8x faster than SAIGE in CPU hours and 26.8x faster in elapsed time. The corresponding carbon footprint estimated using Green Algorithms (see **URLs**) is shown in **Supplement Table S7**, where using REGENIE reduces the CO_2_ footprint by more than 85% compared to SAIGE. **Supplementary Figures S9–S14** compare the accuracy of REGENIE and SAIGE across all 50 traits and show good agreement.

A large portion of the timings for Step 1 of SAIGE is spent fitting the null model following a LOCO scheme, where for each left out chromosome a logistic mixed model is fitted. While the use of the LOCO scheme is well established in the literature^19^ and is used by default in BOLT-LMM^13^ for quantitative traits, it has been suggested that for binary traits it makes little difference^18^. Our experiments suggest it can be very beneficial for binary traits, as demonstrated in **Supplementary Figure S6** where the CAD phenotype is analyzed using SAIGE with and without using LOCO.

### Cross validation scheme

We implemented both a K-fold CV scheme and a leave-one-out CV (LOOCV) scheme in Step 1 for both quantitative and binary traits (see **Methods**). We compared the computational performance and accuracy of both approaches. On both quantitative and binary traits both approaches provide almost identical accuracy (see **Supplementary Figures S15–S16**). On the 50 quantitative traits dataset LOOCV required 192 CPU hours and 23.6GB of memory, whereas K-fold CV required 111 CPU hourse and 12.9GB of memory (see **Table 1**). On the 50 binary traits dataset LOOCV approach requires ~ 50% of the CPU time used by the K-fold CV approach (777 CPU hours versus 1,590 CPU hours) and 65% more memory, but the elapsed time of the two methods is similar (108 hours versus 117 hours). The LOOCV approach requires fewer relatively expensive logistic regression calls compared to K-fold CV, but the extra calls needed are easily parallelized across multiple cores.

### Missing phenotype data

When analyzing multiple traits together with different missing data patterns, we use mean imputation of missing phenotype values in Step 1, but only keep samples with non-missing phenotypes in Step 2. **Supplementary Figures S17, S18 and S19** compares this approximate method to an exact approach that only uses samples with non-missing phenotypes in both Step 1 and Step 2, and shows they are almost identical for both quantitative and binary traits.

### Implementation using Apache Spark

A key advantage of REGENIE is that it can be naturally parallelized over many independent machines using a distributed computing engine such as Apache Spark or Apache Hadoop. By decomposing array SNPs into a block matrix, the subsequent ridge regression and model fitting steps can be attacked with a map-reduce strategy that allows any number of machines to operate independently on single blocks at a time, each of which are of trivial size relative to the full matrix of genotypes. Furthermore, by keeping the block dimensions constant, this strategy allows the method to scale linearly with number of samples or number of SNPs up to arbitrarily large datasets. We implemented a version of this algorithm for quantitative traits in the Glow project (see **URLs**), an open source toolkit for genomic data analysis in Apache Spark. Full details are given in the **Supplementary Methods**. When applied to the same set of 50 UK Biobank quantitative traits as reported in **Table** 1, this approach required 89 hours of CPU uptime and 20.8 minutes of elapsed time for Step 1 when using a Spark cluster of 16 machines, each with 16 cores and 128 GB of RAM, for a total of 256 executors. In terms of elapsed time, this compares favourably to the C++ implementation of REGENIE, which required 12 hours (see **Table** 1).

### Inter-chromosomal LD in the UK Biobank

While developing REGENIE, we identified an anomaly in the UK Biobank array genotypes that led to reduced performance of some of the LMMs being tested. We observed a sizeable number of SNP pairs that exhibited correlation across chromosome boundaries. This issue has been identified independently by other researchers (see **URLs**).

We refer to this as inter-chromosome LD (ICLD). The presence of such SNPs breaks the assumptions of the LOCO scheme used by many LMM and WGR methods, and it can result in loss of power when any one of the SNPs in a pair is associated with a trait. For example, in a test to detect a truly associated SNP that is affected by ICLD, a typical LOCO scheme would remove some or all of that effect when computing phenotypic residuals, as the variant in the pair on the other chromosome would be kept in the null model.

We found that REGENIE can be more robust to ICLD than BOLT-LMM. The effects of SNPs on the trait are first combined within a block and so, if such a pair of ICLD variants were present, the effect of the variants (if large) would be dampened by taking linear combination across the effects of all SNPs in the respective blocks they belong to. As the LOCO scheme in REGENIE is based on the block effects rather than individual SNP effects, REGENIE would then be less prone to a significant decrease in power.

An example of a chromosomal SNP pair found was chromosome 6 and chromosome 15, where a pseudogene *USP8P1* of *USP8* on chromosome 15, a gene involved in cell proliferation, is upstream of *HLA-C* in the major histocompatibility complex (MHC) region, which can harbour large genetic associations for many traits and diseases. In the analysis of LDL with UKB unrelated white British samples, ignoring ICLD resulted in lower strength of association from BOLT-LMM when testing imputed SNPs on chromosome 6 compared to REGENIE. However, once the pairs of SNPs involved in ICLD were excluded from the array SNPs used in step 1, BOLT-LMM resulted in similar p-values as REGENIE (see **Supplementary Figure S20**). A full analysis of ICLD in the UK Biobank data resulted in 3,697 SNP-pairs with significant ICLD (Bonferroni corrected) at the 5% significance level. Many of these pairs are explained due to the presence of gene/pseudogene pairs (see **Supplementary Methods** and **Supplementary Tables S8, S9 and 10** and **Supplementary Figure S21**).

## Discussion

In this study, we present a machine learning method that implements simultaneous whole genome wide regression of multiple quantitative or binary traits. The method uses a strategy that splits computation up into blocks of consecutive SNPs and does not require loading a genome wide set of SNPs into memory. This approach also facilitates analysis of multiple traits in parallel. Overall this results in substantial computational savings in terms of both CPU time and memory usage compared to existing methods such as BOLT-LMM, fastGWA and SAIGE. As the number of large scale cohorts with deep phenotyping grows, this approach will likely become even more relevant. The parallel nature of the approach is ideally suited to distributed environments such as Apache Spark. We have developed a first version of REGENIE for quantitative traits within the Glow project, as well the full version of the method for quantitative and binary traits in a standalone C++ program with source code that is openly available.

One of the biggest computational savings occurs when REGENIE is applied to binary traits and compared to SAIGE. In this setting, we have proposed an approximate Firth regression approach, which we show is almost identical to an exact Firth regression implementation, and much faster. This approach has the added benefit that it avoids the parameter estimate inflation that occurs when SAIGE is used to analyze ultra-rare variants.

The approach used in REGENIE is inspired by, but not the same as, the machine learning approach of stacked regressions^20^. REGENIE uses ridge regression to combine a set of correlated predictors, whereas the Breiman’s stacking approach used non-negative least squares (NNLS) to combine a set of highly correlated predictions in an ensemble learning approach. We have not yet investigated whether NNLS might have advantages here. Additional changes to the method to allow for more flexible priors on the effect size of predictors would also be possible.

There are many potential avenues for development of this approach. It will be easy to expand the functionality to include tests such as SNP × covariate interactions^16^, variance tests^21^ and a whole range of gene-based tests^22–24^. An advantage of analyzing multiple traits simultaneously is that multi-trait tests will be relatively easy to implement in Step 2. Multivariate probit regression for binary traits^25^, multivariate linear regression for quantitative traits^26^ and multi-trait burden tests^27^ will all be straight-forward to implement, with substantial opportunities for application on the cohorts being collected at the Regeneron Genetics Center and around the world.

We also plan to investigate whether REGENIE can be extended to handle time-to-event data^28^ and multinomial regression in a mixed model framework^29,30^, and also the combined analysis of quantitative and binary traits. We suspect it may also be possible to leverage the REGENIE output to estimate SNP heritability, polygenic scores and carry out multi-trait missing data imputation using mixed models on a scale not possible using existing approaches^31^.

Cohorts will continue to grow in terms of sample size, the number of phenotypes and the number of variants available for testing, either via imputation from whole genome sequenced reference panels or via direct whole genome sequencing of study samples. It seems clear to us that Step 1 of the WGR paradigm is now more than computationally tractable using the REGENIE approach. However, further advances will be needed to reduce the computation in Step 2 as whole genome sequencing produces ever increasing numbers of rare variants. These variants can be stored efficiently using the sample indices of rare allele carriers and taking advantage of this can substantially reduce the cost of computation of sums and products needed for test statistic calculations.

## Online Methods

### Whole genome linear regression

In a sample of *N* individuals, let **y** denote the *N*-element phenotype vector, **G** represent the *N* × *M* genotype matrix, where *G_ij_* ∈ {0, 1, 2} is the allele count for individual *i* at the *j*-th marker, and **X** represent the *N* × *C* matrix of covariates (including an intercept) which is assumed to be full rank. We consider a whole genome regression model

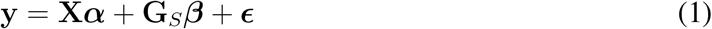

*α* are the fixed covariate effects, **G**_*S*_ is a standardized version of **G** where the genotypes have been transformed to have mean 0 and variance 1, 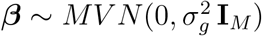, and 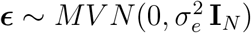. This is the standard infinitesimal model, which can also be re-written as

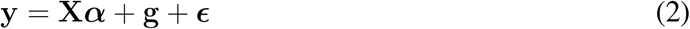

with 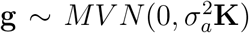, where 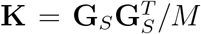 is usually referred to as a genetic relatedness matrix (GRM) or empirical kinship matrix, and 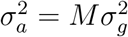 is the additive polygenic variance.

Covariates effects are removed from both the trait and the genotypes in Eq(1) by first computing an orthonormal basis for the covariates, projecting the genotypes and the trait onto that basis, and then subtracting out the resulting vectors to obtain the residuals. This is equivalent to using a projection matrix **P**_*X*_ = **I**_*N*_ – **X**(**X**^*T*^**X**)^−1^**X**^*T*^ with

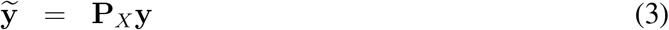

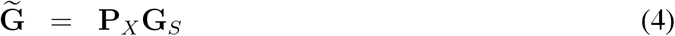

Both the genotype and phenotype residuals are then scaled to have variance 1.

### Stacked block ridge regression

Fitting Eq(1) is computationally intensive since **G** has typically many hundreds of thousands of columns. Instead, for Step 1, we transform the model to

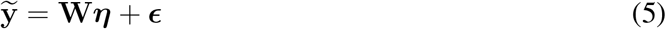

where **W** is a matrix derived from **G** with many fewer columns. Specifically, we divide **G** into blocks of *B* consecutive and non-overlapping SNPs, and from each block we derive a small set of predictors, using ridge regression across a range of *J* shrinkage parameters (see **Supplementary Methods**). The idea behind using a range of shrinkage values is to capture the unknown number and size of truly associated genetic markers within each window. This approach is equivalent to placing a Gaussian prior on the effect sizes of the SNPs in the block and finding the maximum a posteriori (MAP) estimate of the effect sizes and the resulting prediction. Another approach would be to integrate out the effect sizes over the Gaussian prior to obtain the Best Linear Unbiased Prediction (BLUP)^32^, but we have not investigated that approach in this paper.

The ridge predictors are re-scaled to have unit variance and are stored in place of the genetic markers in matrix **W**, providing a large reduction in data size. If, M=500,000 and B=1,000 and J=5 shrinkage parameters are used, then the reduced dataset will have JM/B = 2,500 predictors. We refer to this part of the method as the *level 0* ridge regression.

In order to keep memory usage low when analyzing multiple traits, the within-block predictions are stored on disk and read separately for each trait when fitting models at *level 1* (see below). The added I/O operations incur a small cost on the overall runtime and significantly decreases the amount of memory needed.

The ridge regression takes account of linkage disequilibrium (LD) within each block, but not between blocks. One option that we have considered, but not implemented yet, is to condition the ridge regression on the estimates from the previous block, and this may better account for LD across block boundaries.

The predictors in **W** will all be positively correlated with the phenotype. Thus, it is important to account for that correlation when building a whole genome wide regression model. The predictors will also be correlated with each other, especially within each block, but also between blocks that are close together due to LD. We use a second level of ridge regression on **W** for a range of shrinkage parameters and choose a single best value using K-fold cross validation scheme^20^. This assesses the predictive performance of the model using held out sets of data, and aims to control any over-fitting induced by using the first level of ridge regression to derive the predictors (see **Supplementary Methods**). We refer to this part of the method as the *level 1* ridge regression.

The result of this model fit is a single *N* × 1 predicted phenotype 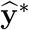, and this can be partitioned into 22 leave-one-chromosome-out (LOCO) predictions (denoted 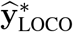) which are used when testing SNPs for association in Step 2 to avoid proximal contamination (see **Supplementary Methods**).

### Association testing

In Step 2, when testing for association of the phenotype with a variant **g**, we consider a simple linear model

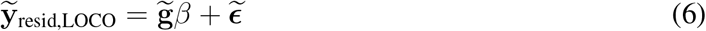

where 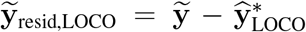, refers to the phenotype residuals where the polygenic effects estimated from the null model with LOCO have been removed, 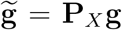 are residuals obtained from removing covariate effects from the tested variant, and 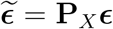 with 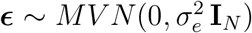. A score test statistic for *H*_0_: *β* = 0 is

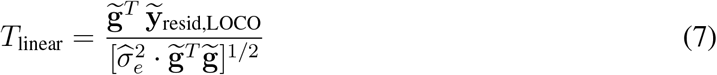

where we use 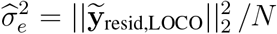 (note that in large scale applications, *N* is very large so that *N* – *C* ≈ *N*). In Eq(7), when estimating the variance of the term in the numerator, we assume that the polygenic effects are given which leads to the denominator involving only *O*(*N*) computation. While other methods make use of a calibration factor in the denominator to account for the variance of the polygenic effects^13,18,33^, we found in applications that the results obtained using this simple form match up closely to those using a calibration factor. Finally, we use a normal approximation 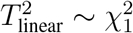 to estimate the p-value. As with Step 1 above, the REGENIE software reads the genetic data file in blocks of *B* SNPs, and these are processed together, taking advantage of parallel linear algebra routines in the Eigen library.

### Multiple traits

Both Step 1 and Step 2 above are easily extended so that multiple phenotypes can be processed in parallel. The genetic data files in both steps can be read once, in blocks of *B* SNPs, which means the method uses a small amount of memory. In addition, the linear algebra operations for the covariate residualization, ridge regression and association testing can be shared across traits. This is similar to the approach implemented in the BGENIE software (see **URLs**) for single SNP linear regression analysis^17^. Fine details of the multiple phenotype approach are given in the **Supplementary Methods**.

### Binary traits

For binary traits we use exactly the same level 0 ridge regression approach, which effectively treats the trait as if it were quantitative. However, at level 1, instead of a linear regression in Eq(5) we use logistic regression

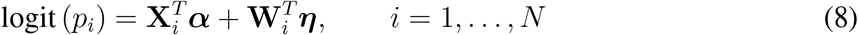

where 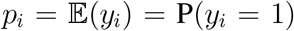 with *y_i_* indicating the case status of the *i*-th individual, **X**_*i*_ is the covariate vector for the *i*-th inidividual, *α* are the fixed covariate effects, **W**_*i*_ are the (*BR*) within-block predictions for the *i*-th individual, and ***η*** = (*η*_1_,…, *η_BR_*)^*T*^ with 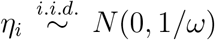. This model corresponds to logistic regression with ridge penalty applied to the effects of within-block predictions in **W**.

We approximate the model in Eq(8), by first fitting a null model for each trait that only has covariate effects

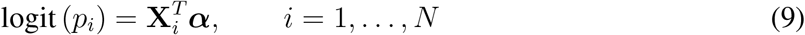

and then using the resulting estimated effects as an offset in the model in Eq(8),

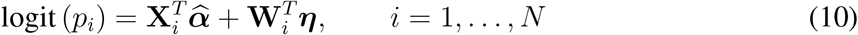

where 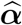 represent the effects estimated in Eq(9). As covariate effects are not expected to strongly change (unless correlation between covariates and block predictions are very large), this approximation is expected to work well in most analyses.

As with quantitative traits, we use *K*-fold cross validation to choose the level 1 ridge regression parameter. However, for extremely unbalanced traits, it may happen that one of the folds contains no cases. To avoid this situation we also implemented an efficient version of leave one out cross validation (LOOCV). While at first sight it may seem that LOOCV is more computationally intensive than *K*-fold CV since the model has to be fitted *N* times (rather than *K* times) on data with (*N* – 1) samples, the leave-one out (LOO) estimates can actually be obtained (approximately for binary traits) from rank 1 updates to the results from fitting the model to the full data once (see **Supplementary Methods**). We have found in practice that LOOCV gives similar association results as *K*-fold cross validation (see **Supplementary Figures S15** and **S16**) and in some cases can be computationally faster (see **Tables 1–2**). A LOCO scheme is applied to the polygenic effect estimates and the resulting predictions 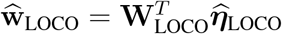 are then stored.

In Step 2, we use a logistic regression model score test to test for association between each marker and binary trait. Covariate effect sizes are estimated along with genetic marker effect sizes, but we include the LOCO predictions from Step 1 as a fixed offset (see **Supplementary Methods**).

When rare variants are tested for association with a highly unbalanced traits (i.e. a trait that has low sample prevalence), the use of asymptotic test statistic distributions does not work well, and results in elevated Type I error rates. REGENIE implements several methods to handle this situation. Firstly, it includes the saddle pointapproximation (SPA) test^34^ that is also included in SAIGE^18^. This approach better approximates the null distribution of the test statistic, but we have found that it can sometimes fail to produce good estimates of SNP effect sizes and standard errors, which are higly desirable for meta-analysis applications (see **Supplementary Table S4** and **Supplementary Figure S5**).

Secondly, we use Firth logistic regression, which uses a penalized likelihood to remove much of the bias from the maximum likelihood estimates in the logistic regression model. This approach results in well calibrated Type 1 error and usable SNP effect sizes and standard errors. Since the use of Firth regression can be relatively computationally intensive, we have developed an approximate Firth regression approach that is much faster (**Supplementary Table S5**), which involves estimating covariate effects in a null Firth regression model, and then including covariate effects along with the LOCO genetic predictor as offset terms in a Firth logistic regression test (see **Supplementary Methods**). In practice, we have found this approximation to give very similar results to using the exact Firth test (**Supplementary Figure S7**).

### Handling missing data

As a key goal of our approach is to analyze multiple traits all at once, one issue that remains to be addressed is the presence of missingness in the data, which could differ among the traits. We consider different approaches based on the nature of the trait as well as whether the null model is being fitted or whether association testing is being performed.

For quantitative traits, missing data when fitting the null model is addressed by replacing missing values by the sample averages for each trait. In the association testing step, in addition to the option of mean-imputing missing entries in the phenotypes, we also consider an alternative where for each trait, individuals with missing observations are removed from the analysis. This is done by ensuring that when taking sums over individuals, those with missing phenotype have a zero contribution to the sum. This is similar to the approach implemented in the BGENIE software (see **URLs**) for single SNP linear regression analysis^17^. We assume that covariates are fairly well-balanced in the sample and project them out of the phenotypes using all the samples ignoring the missingness within each trait. In the case where phenotypes have the same or very similar patterns of missingness, or if only a single phenotype is being analyzed, it may be more logical to discard missing observations rather than impute them with the sample averages per trait. Hence, we implement an alternative approach where in both the null fitting step and the association testing step, all samples with missingness at any of the *P* phenotypes are dropped. An approach we have not yet implemented, but may produce better results for quantitative traits would involve using a multivariate normal model to jointly model correlation between the set of traits and impute missing data, either before, or conditional upon the output of Step 1.

For binary traits, we use the mean-imputed phenotypes to fit the level 0 linear ridge regression models within blocks, but discard missing observations when fitting level 1 logistic ridge regression. As the logistic ridge regressions are fitted separately for each trait, this makes it straightforward to account for the missingness patterns separately for each trait. Similarly, in the testing step, we discard missing observations when fitting logistic regression separately for each trait as well as when using Firth or SPA corrections.

### UK Biobank data set

The UK Biobank^17^ is a large prospective study of about 500,000 individuals between 40-69 years old with extensive phenotype information being recorded. Genotyping was performed using the Affymetrix UK BiLEVE Axiom array on an initial set of 50,000 participants, and the Affymetrix UK Biobank Axiom array on the remaining participants. Up to 11,914,699 variants imputed by the Haplotype Reference Consortium (HRC) panel that either have minor allele frequency above 0.5% or have minor allele count above 5 and are annotated as functional in 462,428 samples of European ancestry were used in the data analyses. We selected up to 407,746 individuals of white British ancestry who had genotype and imputed data available and applied quality control filters on the genotype data using PLINK2^35^, which included MAF ≥ 1%, Hardy-Weinberg equilibrium test not exceeding 10^−15^ significance, genotyping rate above 99%, and LD pruning using a R^2^ threshold of 0.9 with a window size of 1000 markers and a step size of 100 markers. This resulted in up to 471,762 genotyped SNPs that were kept in the analyses.

### Statistical analyses

We used REGENIE to perform GWA analyses on up to approximately 11 million imputed variants for 50 quantitative traits and 54 binary traits of up to 407,746 white British participants in the UK Biobank. Quantitative phenotypes were converted to z-scores using rank-inverse based normal transformation (RINT). In the statistical models used, covariates included age, age^2^, sex, age× sex and the top 10 principal components provided by the UK Biobank. To assess the performance of REGENIE in GWAS, we compared the results from REGENIE to those of existing approaches for large-scale analysis, which included BOLT-LMM (version 2.3) and fastGWA (GCTA version 1.93.0beta) for quantitative phenotypes, and SAIGE (version 0.36.5.1) with the LOCO option for binary traits. For all methods, Step 1 was run on a set of array SNPs stored in bed/bim/fam format and Step 2 was run on imputed data stored in the BGEN format. All of the programs were called from within R^36^, where we used the function system.time to keep track of CPU and wall clock timings for each run.

## Data availability

The individual-level genotype and phenotype data are available through formal application to the UK Biobank http://www.ukbiobank.ac.uk.

## Code availability

The C++ source code for REGENIE is available from https://rgcgithub.github.io/regenie/ under an MIT License.

## URLs

REGENIE https://rgcgithub.github.io/regenie/

BGENIE https://jmarchini.org/BGENIE/

SNPTEST https://mathgen.stats.ox.ac.uk/genetics_software/snptest/snptest.html

UK Biobank http://www.ukbiobank.ac.uk

PLINK https://www.cog-genomics.org/plink2

BOLT-LMM (version 2.3) https://data.broadinstitute.org/alkesgroup/BOLT-LMM/

SAIGE (version 0.36.5.1) https://github.com/weizhouUMICH/SAIGE

fastGWA (GCTA version 1.93.0beta) https://cnsgenomics.com/software/gcta/#Overview

Glow http://projectglow.io

LDstore (version 2.0beta) http://www.christianbenner.com/

Green Algorithms http://www.green-algorithms.org

## Acknowledgments

We are grateful to Frank Austin Nothaft, Henry Davidge, Kiavash Kianfar, Karen Feng and Yong-sheng Huang for ongoing advice on developing the REGENIE code within the Databricks environment.

## Author contributions

J.Ma conceived and supervised the study. J.Ma, J.Mb, L.B and E.M developed the method for quantitative traits. J.Mb and J.Ma developed the method for binary traits. J.Mb and J.Ma coded the C++ implementation of the method. J.Mb carried out all the testing and real data analysis of the C++ method. L.B and E.M developed the Apache Spark implementation of the method. C.B provided advice and code for LD calculations. J.B, A.M, J.K tested and provided comments on the C++ version. J.Ma and J.Mb wrote the manuscript. All other authors provided comments at various stages of the project.

## Supplementary Methods

### Block partitions

The genotype matrix 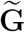 is partitioned into blocks, and only a single block is read into in memory at a time. Let 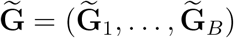 denote a partition of 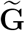 into *B* contiguous blocks of SNPs, where all SNPs within the same block are from the same chromosome. Many choices are available to determine how to define the blocks (e.g. physical or genetic distance, using LD information). In practice, we fix the the block size to be the same (approximately) across blocks. Let *m_i_* denote the number of SNPs within the *i*-th block, so that 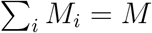.

### Level 0 ridge regressions

Given a genotype block 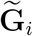 of *M_i_* SNPs, we aim to reduce the number of predictors in the block *M_i_* to a much smaller number which would capture both the LD that may exist between the markers in the block as well as any effects they may have on the trait. To address the former, we consider fitting a penalized linear model with a *L*2 penalty (known as ridge regression^37,38^)

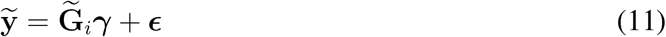

where ***ϵ*** = (*ϵ*_1_,…, *ϵ_N_*)^*T*^ with 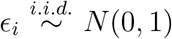, and we obtain an estimate for the marker effects *γ* as

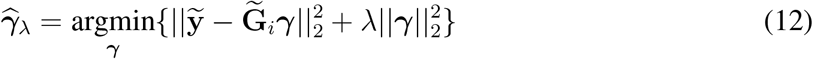

The parameter λ in Eq(12) is often referred to as the shrinkage parameter, as higher values induce more shrinkage towards zero in the effect size parameter *γ* and lower values of λ reflect larger effect sizes. The estimate 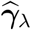 can be viewed from a Bayesian framework as the maximum a posteriori (MAP) estimator in the model in Eq(11) where a Gaussian prior is used for the marker effect sizes

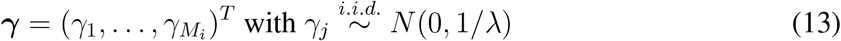

Given these model assumptions, we can express the ridge penalty parameter as a function of the SNP-heritability 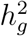, where we have 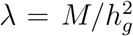. As we do not know what the true effects of the markers on the trait are, we consider a set of parameter values Λ = {λ_1_,…, λ_*R*_}, which aim to reflect a comprehensive range of values for the SNP-heritability of the trait. More precisely, we choose *R* evenly spaced values in [0, 1] for 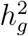, where the minimum and maximum values are set to 0.01 and 0.99, respectively, and compute the corresponding λ parameter values given the total number of SNPs *M*. The solutions from ridge in Eq(12) can be computed analytically with the closed-form expression

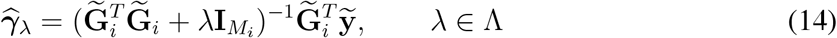

and so for each block of size *M_i_*, we end up with a much smaller set of *R* predictors,

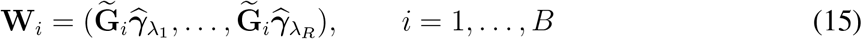

Let **W** = (**W**_1_,…, **W**_*B*_) be the resulting *N* × (*BR*) matrix obtained from fitting the ridge regressions on all the *B* genotype blocks. Assuming that the block size and the number of ridge parameters is kept reasonably small, (*BR*) should be much smaller than the number of markers *M*, so that memory usage would be much lower than when reading the whole genotype matrix at once. For example, in a sample of *N* = 500, 000 individuals with *M* = 500, 000 markers and a block size of *T* = 1, 000 with *R* = 5 ridge parameters, using this approach would save more than 52GB in memory usage (assuming that loading the genotype matrix costs MN/4 bytes of memory) and the number of predictors would be reduced to *BR* = 2,500.

### Level 1 ridge regression

To combine the predictors in **W**, we consider an approach commonly used in the statistical and machine learning literature, referred to as stacking^20^, where the aim is to obtain an optimal linear combination of the block predictions using penalized regression. This approach has been shown to lead to better predictions than using a single predictive model to fit the data. We re-scale the predictors in **W** to have unit variance and apply ridge regression, which corresponds to the following model

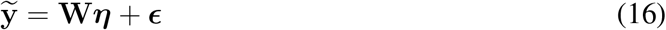

where ***ϵ*** = (*ϵ*_1_,…, *ϵ_N_*)^*T*^ with 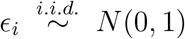 and solutions for ***η*** are computed by adding a *L*2 penalty to the log-likelihood function of the model in Eq(16) which results in the following estimator

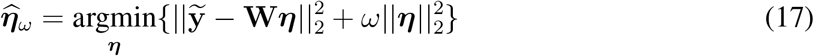

Similarly to Eq(14), it has a closed-form expression

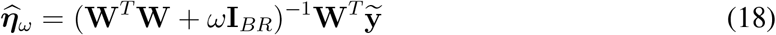

The hyper-parameter *ω* is selected from a set of *Q* shrinkage parameters Ω = {*ω*_1_,…, *ω_Q_*}, where given an assumed SNP-heritability value 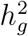, we express the corresponding ridge penalty parameter as 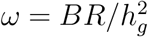. An optimal value *ω*^*^ is chosen by minimizing the the prediction error

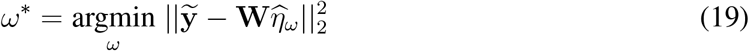

### *K*-fold cross-validation

As the block predictions **W** are correlated with the phenotype, *K*-fold cross-validation (CV) is used to reduce over-fitting. More precisely, we split the data into *K* folds and using the subscript (.)_(−*k*)_ to represent the data where the *k*-th fold has been removed and (.)_(*k*)_ to represent the data for the *k*-th fold, we obtain out-of-sample block predictions for each fold *k* by fitting the model in Eq(11) in each block **G**_*i*_ leaving out fold *k*

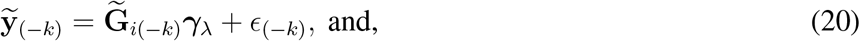

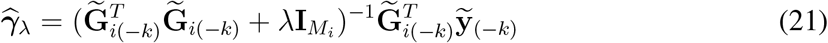

Hence, out-of-sample predictions for fold *k* in the *i*-th genotype block are obtained as

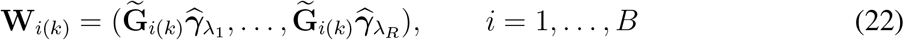

This is done within each block 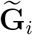 across all *K* folds. Let **W**^*^ denote the *N* × (*BR*) matrix containing the out-of-sample predictions across the *B* blocks. We obtain the prediction error for fold *k* by leaving out the fold when fitting the model in Eq(16) and using the resulting estimates to obtain the predictions

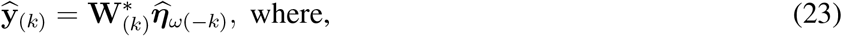

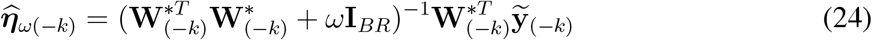

We select the value of *ω* in Eq(24) that minimizes the prediction errors over the *K* folds

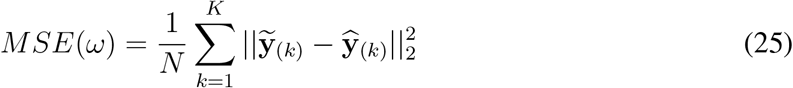

### LOCO predictions

Let 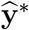 refer to the final predictions obtained from Eq(24) by using the optimal hyper-parameter chosen from CV over all the K folds. Instead of using 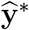 which captures the polygenic effects genome-wide, we use an approximate LOCO scheme so that the resulting predictions only capture effects outside of the chromosome containing the SNP being tested. More precisely, for each chromosome, we compute the whole-genome predictions by setting the entries in 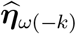 in Eq(23) corresponding to blocks in the chromosome to 0. This approach is less computationally burdensome than a more exact LOCO scheme which would consist in leaving out all SNPs within the tested chromosome before partitioning the genome into blocks. We have found that it works well in practice.

### Multiple traits

In the context of multiple traits, we simply replace the *N* × 1 phenotype vector **y** by an *N* × *P* matrix of *P* phenotypes when projecting out covariates in Eq(3) to obtain a matrix of phenotypic residuals 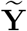. We then use this matrix in Eq(14) to obtain predictions for all traits at once within each genotype block. This is one of the key steps of the method as the genotype data only has to be read once for all the traits and many of the operations involved on the genotype block are the same for all traits. Hence, it highly reduces the computational burden involved in the null model fitting step when many traits are analyzed. Since the block predictions will differ across traits, we then fit the model in Eq(16) separately for each trait. This increases the computational burden in memory usage, as a *N* × (*BR*) matrix has to be stored separately for each of the *P* traits (in our example above with *N* = 500, 000 individuals, *M* = 500, 000 markers, *T* = 1, 000 markers in a block and *R* = 5 ridge parameters, this would correspond to about 10GB of memory used per trait). To address this, the block predictions can be stored on disk as each genotype block is read in memory so that the memory usage when going through blocks is kept relatively low. Then, when fitting the model in Eq(16) for each trait, only the block predictions for the trait being analyzed need to be stored in memory, which reduces the memory usage to the same as that used when analyzing a single trait.

### Whole genome logistic regression for binary traits

For binary traits, a logistic regression model is used at level 1. This model handles covariates by estimating their effects in a null model (Eq(9)) that only includes covariates, and these are then used as an offset term in a full logistic regression model (Eq(8)). The Newton-Raphson algorithm is used to obtain parameter estimates in Eq(9) and Eq(10). To obtain the optimal parameter from *K*-fold CV, we use the average log-likelihood as the loss function to measure performance

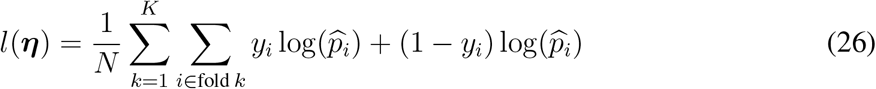

where for individual *i* in fold k, 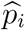 is the computed from Eq(10) using the estimate for ***η*** which was obtained by fitting the model on the remaining (*K* – 1) folds. The covariate effects are not re-estimated within each fold as we assume they should be fairly well-balanced across the *K* folds. The optimal parameter chosen for ***η*** is the one that minimizes – *l*(***η***).

As with quantitative traits, a LOCO scheme is applied to the polygenic effect estimates and the resulting predictions 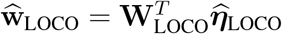 are then stored for step 2.

### Logistic regression for case-control association tests

To test a variant **g** = (*g*_1_,…, *g_N_*)^*T*^ for association with a single binary trait, we consider the following model

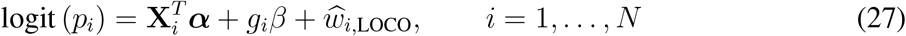

where the polygenic effects predictions 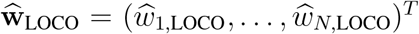 stored from step 1 are included as an offset in the logistic model. A score test statistic for *H*_o_: *β* = 0 is

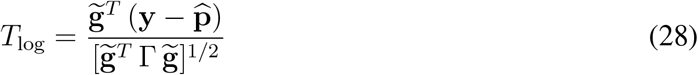

where 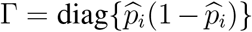 with 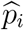 being the estimated mean of the trait under the null hypothesis for the *i*-th individual, and 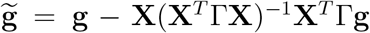 is the genotype residual vector after adjusting for covariates. Similar to with quantitative traits, no calibration factor is used in the test statistic to account for the variance in the polygenic effects estimate. The p-value is estimated using a normal approximation with 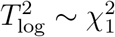.

### Firth logistic regression

When rare variants are tested for association with a highly unbalanced traits (i.e. a trait that has low sample prevalence), quasi-complete separation can occur in logistic regression where none of the cases contain the minor allele which leads to very unstable estimates of the effect sizes and standard errors. One solution to address this is with Firth correction^39,40^, where a penalty based on the observed Fisher information matrix is added in the log-likelihood

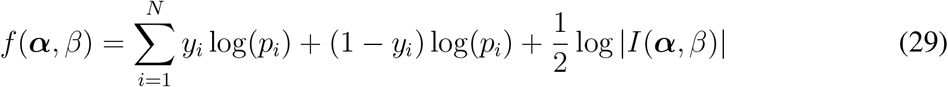

where *I*(***α**, β*) = **U**^*T*^Γ**U** with **U** = (**X g**), and Γ = diag{*p_i_* (1 – *p_i_*)} where the linear predictor is the same as in Eq(27). This penalized likelihood function is equivalent to the posterior distribution function using Jeffreys invariant prior^39^. The last term in Eq(29) is maximized when *p_i_* = 0.5 which occurs when ***α*** = **0** and *β* = 0, and hence, adding this penalty shrinks the coefficients towards 0. When rare variants are tested with highly unbalanced traits, the literature suggests^40^ that a Wald test within the Firth regression approach may not work well. A score test may be a possibly fast solution that we implement in future versions, but in this first version we use a likelihood ratio test (LRT) to test the null hypothesis *H*_0_: *β* = 0

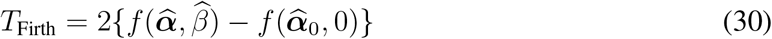

where 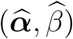 is the maximum of the penalized likelihood function and 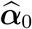 is the maximum of the penalized likelihood when *β* = 0. The p-value is estimated using 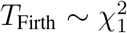. As the penalty term in Eq(29) involves the tested variant **g**, the second term in Eq(30) has to be re-evaluated for each tested variant, which means the penalized likelihood Eq(29) with *β* = 0 has to be maximized for each tested variant. This can render the test very computationally burdensome, all the more when many covariates are included as it determines the dimension of the matrix involved in the penalty term in Eq(29).

### Approximate Firth logistic regression

To ease the computational burden, we also consider an approximation to the test in Eq(30), where we first estimate covariate effects in a null model where we include a modified penalty term,

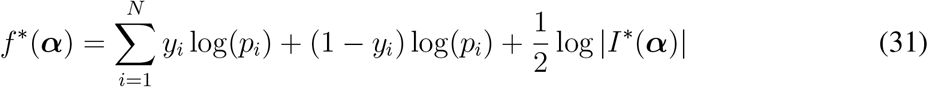

where *I*^*^(***α***) = **X**^*T*^Γ**X** with Γ = diag{*p_i_*(1 – *p_i_*)}, and the linear predictor is the same as in Eq(27) with *β* = 0. The penalty term in *f*^*^ does not depend on the tested variant which means that *f*^*^ only needs to be maximized once per chromosome tested to obtain an estimate for the covariate effects. Let 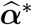 denote the resulting estimate of the covariate effects. We consider the modified test statistic

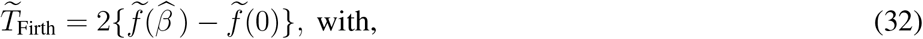

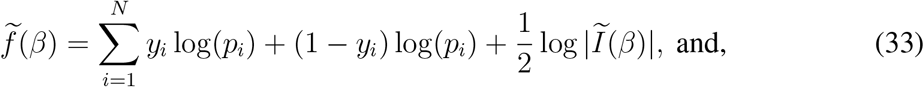

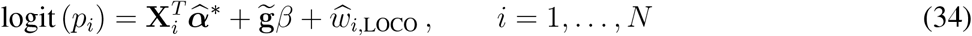

where 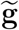 are residuals obtained from removing covariate effects (i.e. 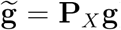), and the observed Fisher information 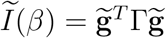 is a scalar which means the computations due to the penalty term are *O*(1) rather than *O*(*K* + 1). The mean function in Eq(34) incorporates covariate effects through an offset term, which reduces the number of predictors from *K* + 1 down to a single predictor, the tested variant. In practice, we have found this approximation to give very similar results to using the exact Firth LRT.

### Saddle Point Approximation (SPA) test

In the same context where rare variants are being tested, several methods have proposed using a saddle pointapproximation (SPA) rather than a normal approximation to evaluate significance of the test in Eq(28) and found that it resulted in better calibration of the test statistic when traits were highly unbalanced^18,34^. Rather than relying on the first two moments of the test statistic to approximate its null distribution, SPA approximates the cumulant generating function of the test statistic, which involves all of the higher order moments. Hence, SPA can lead to better approximations in the tail of the null distribution compared to the normal approximation. For the test statistic in Eq(28), which is a linear combination of Bernoulli random variables, the cumulant generating function (CGF) is

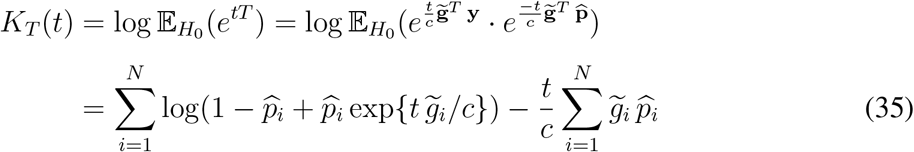

where *T* = *T*_log_, and 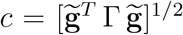 is the denominator of the test statistic. The first and second derivative of *K_T_*(*t*) can also be obtained from Eq(35) as

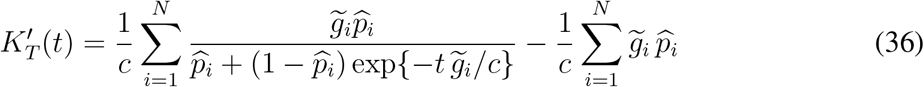

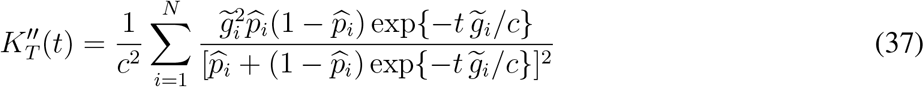

Given an observed test statistic value *t*_obs_, the distribution function of *T* is obtained using a saddle pointapproximation^41^ as

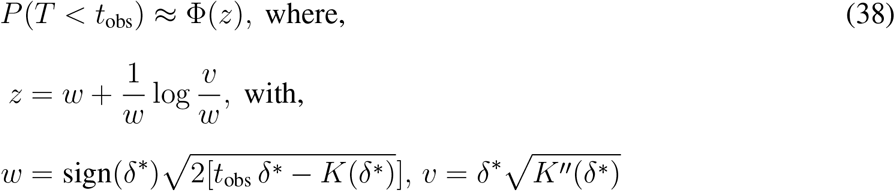

where Φ denotes the standard normal distribution function and *δ*^*^ is obtained by solving the equation *K′*(*t*) = *t*_obs_, which we do using a modified Newton-Raphson algorithm that uses the bisection method. To avoid having to solve the equation twice for both tails of the distribution (as the statistical test performed is two-sided), we use the fact that *K*_−*T*_(*t*) = *K_T_*(–*t*) so that we only need to approximate one of the tails of the null distribution. The SPA approximation can fail for extremely rare variants (e.g. minor allele count [MAC] less than 5) so we use a minimum MAC filter in our implementation where the default is set to 5. Similarly to previous methods^18,34^, we also make use of the sparsity in the genotype vector when the MAC is low by approximating the CGF *K_T_* and its derivatives using a normal approximation in order to obtain functions which only involve sums over the non-zero entries of the genotype vector rather than sums over all of the entries.

### Short derivation of the SPA approach

We have found the description of the SPA approximation in the statistical genetics literature to be rather opaque^18,34^, so here we provide a brief derivation of the result stated in Eq(38), based on the material in Butler (2007).

Suppose *X* is a continuous random variable with probability density function (pdf) *f*(*x*), 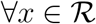. The moment generating function (mgf) of *X* is defined as

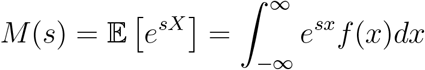

over all values of *s* for which the integral converges.

The cumulant generating function (cgf) of *X* is defined as

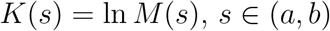

where (*a, b*) is the largest interval of convergence. Some basic properties of the mgf and cgf are 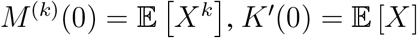.

The mgf can be re-expressed as

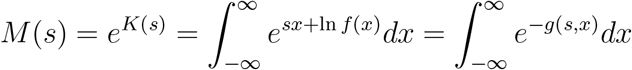

where *g*(*s, x*) = –*sx* – ln *f*(*x*), and then approximated using Laplace’s method as

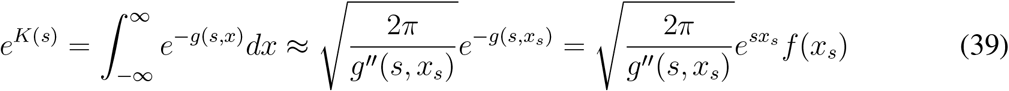

where *x_s_* = argmin *g*(*s, x*) is the solution of

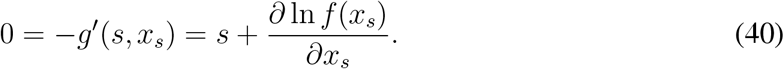

At this point, it is useful to recognize that *x_s_* is a function of *s* and vice-versa.

Eq(39) can be expressed as

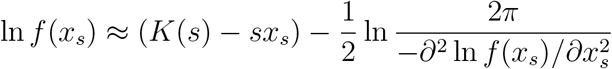

and assuming the second term is constant with respect to *x_s_*, we have

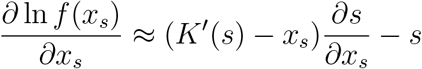

and then Eq(40) becomes

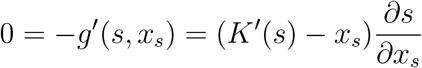

so the definition of *x_s_* reduces to *x_s_* = *K′*(*s*). We also have

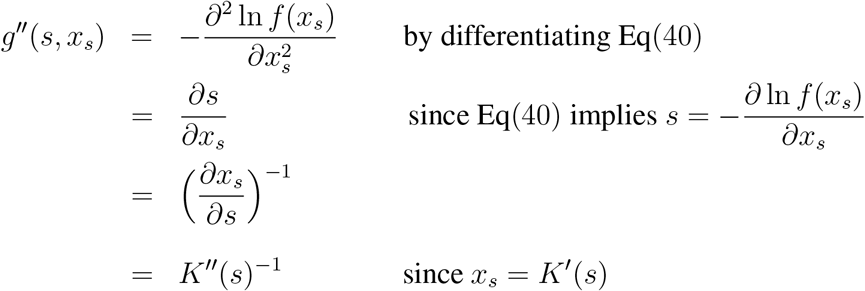

The saddle pointapproximation to the pdf of *X* is then the re-expression of Eq(39) as

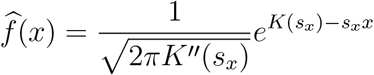

where *s_x_* is a function of *x* that satisfies *x* = *K′*(*s_x_*).

The SPA to the cumulative density function (cdf) of *X*, defined as *F*(*y*) = *P*(*X* < *y*) can be approximated using the SPA of the pdf of *X* as

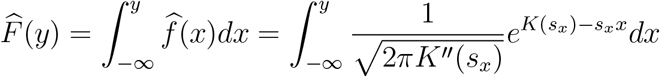

Then define 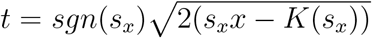, which is a function of *x*. This means the integral can be re-expressed to involve the standard normal density function *ϕ*(.) as

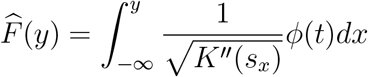

It can be shown^41^ that

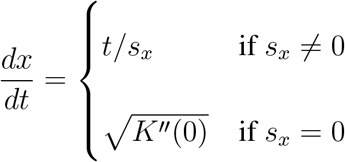

and changing variables from x to t results in

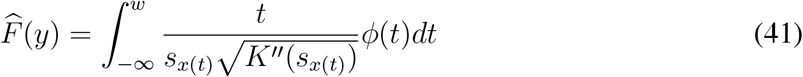

where *s*_*x*(*t*)_ is *s_x_* expressed as a function of *t*, 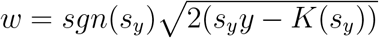 is a function of *y* and *K′*(*s_y_*) = *y*.

We then use the Temme approximation, which states that if *ϕ*(.) and Φ(.) are the standard normal pdf and cdf respectively, then

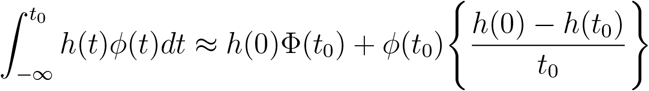

Applying this to Eq(41) using 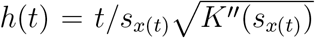 and *t*_0_ = *w*, for which it can be shown that *h*(0) = 1, and defining 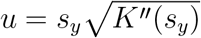, we get

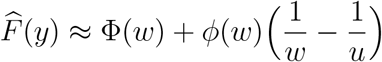

Finally, defining 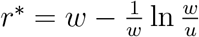 results in the final SPA shown in Eq(38)

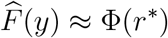

where the approximation occurs via a first order Taylor approximation of both Φ(.) and ln(.).

### Leave-one out cross-validation (LOOCV)

As an alternative to *K*-fold CV, we implement LOOCV which is equivalent to performing *N*-fold CV. With binary traits, LOOCV can be very useful when the number of cases in the sample is low, which in *K*-fold CV can result in a fold that only contains controls. While at first sight it may seem that LOOCV is more computationally intensive than *K*-fold CV since the model has to be fitted *N* times (rather than *K* times) on data with (*N* – 1) samples, the leave-one out (LOO) estimates can actually be obtained (approximately for binary traits) from rank 1 updates to the results from fitting the model to the full data once. For example, with a quantitative trait where we consider the following ridge regression model

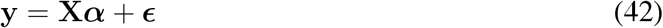

where **X** is a *N* × *C* matrix of predictors and a *L*2 penalty is applied to the parameter ***α***, the LOO estimate for the *i*-th individual is

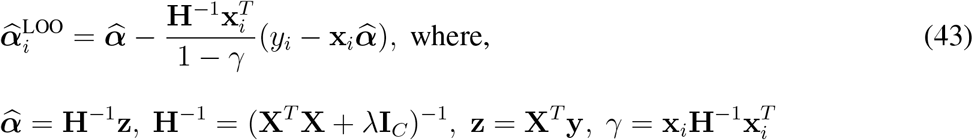

with 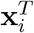 representing the *C* × 1 vector of predictors for the *i*-th individual and λ being the ridge penalty parameter. Both **H**^−1^ and **z** in Eq(43) are already computed when fitting the model on the full data set and 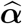 is the resulting estimate of that fit. The remaining operations needed to obtain the LOO estimate only involve at most matrix-vector operations. We perform these operations for groups of individuals at a time in order to have matrix-matrix operations which use multiple threads in the Eigen C++ library. Also, when considering a range of penalty parameters, many of the operations in Eq(43) are the same, and so we only compute them once and re-use them for each penalty parameter.

For binary traits, we consider a logistic ridge model where the penalized log-likelihood function is

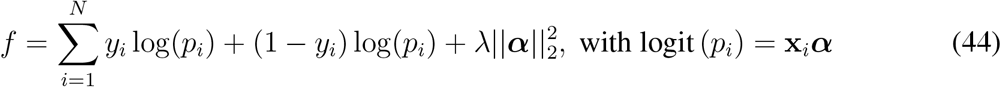

and *p_i_* is the marginal phenotypic mean for the *i*-th individual. The parameter estimate 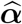 is then obtained by maximizing the function in Eq(44) using the Newton-Raphson algorithm starting at the origin. Assuming that the estimate obtained after removing a single observation should be fairly close to that obtained from the full data fit, we approximate the LOO estimate for individual *i* by solving the function in Eq(44) where individual i has been excluded, and where we start at the solution 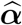 instead of starting at the origin, and we perform only a single iteration of the algorithm. The resulting parameter estimate can be expressed in closed-form

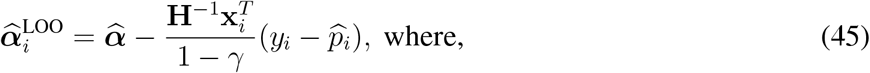

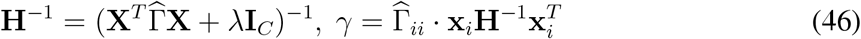

and 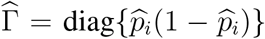 with 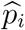 being the estimated mean from the whole data fit. The matrix **H**^−1^ is also computed from that same model fit so we only need to perform at most matrix-vector operations, for 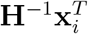, which can be done for groups of individuals at a time to have matrix-matrix operations. The remaining vector-vector operation 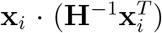 needed is *O*(*C*). Using these computational tricks, we are able to efficiently obtain LOO estimates based on a ridge regression model for both quantitative and binary traits.

### Time complexity of REGENIE

**Supplementary Table S1** gives an overview of the computational complexity of the REGENIE method. It can be broken down based on the two main steps, which are whether the null model is being fitted (step 1), or whether variants are being tested for association (step 2). In step 1, two levels of models are considered which correspond to linear mixed models applied within genotype blocks (level 0) and across all genotype blocks (level 1). By leveraging the fact that many of the operations done within a block are the same for all the phenotypes, we are able to reduce the time complexity involved in the level 0 models where the operations done once for all traits have time complexity of about *O*(*B*^3^*J*_0_) times the total number of blocks. This is due to the spectral decomposition of the genetic correlation matrix within each block, which only needs to be computed once with a time complexity about *O*(*NB*^2^), where both the number of SNPs within a block *B* and the number of level 0 ridge parameters *J*_0_ are kept relatively small (e.g. *B* = 1,000, *J*_0_ = 5). Similarly, the computation time for the level 1 models is driven by the spectral decomposition of the correlation matrix based on the *R* level 0 within-block predictions, where *R* is still relatively small compared to the total number of SNPs in step 1 (e.g. *R* = 2,500 for *M*_1_ = 500,000). Hence, step 1 of REGENIE can lead to better computational efficiency relative to BOLT-LMM and SAIGE which are *O*(*N*^1,5^ *M*) times the number of phenotypes, where in large scale applications *N*, in addition to *M*, would be large (e.g., *N* = 500,000). For step 2, all three methods have the same order of complexity which is *O*(*M*_2_*N*) times the number of phenotypes, where *M*_2_ is the number of variants being tested for associations.

### Optimization of the association testing step in REGENIE

Step 2 of REGENIE can become a major computational bottleneck when millions of SNPs have to be tested for association with multiple traits. Hence, we optimized this step when the genotype file input is BGEN v1.2 format with 8-bit encoding, which is the file format used by UK Biobank for the tens of millions of imputed SNPs. More precisely, REGENIE has been optimized to read the BGEN file with multiple threads by taking advantage of the highly structured format of BGEN files. We initially do a first pass though the BGEN file to get the index of the markers in the file while skipping over the genotype data. We process the tested SNPs in blocks and for each block, we first open the file using the index information of the first variant in the block and then read the compressed genotype data for each variant. We decompress the genotype data, impute missing data, remove covariate effects, scale the residuals and compute the association test statistics for each variant. We experimented using the multi-threading in Eigen to efficiently parallelize as many matrix × matrix operations as possible. However, we found that a better strategy was to disable this feature and instead use OpenMp to parallelize the computations over the markers.

### Details of Apache Spark implementation

The following procedure describes the map-reduce strategy used in Glow (see **URLs**) for transforming the standardized genotype matrix *G* into the matrix of ridge predictors W for a set of quantitative phenotypes *Y* using a set of ridge regularization parameters *J*. Let each block *g* in matrix *G* be indexed by a sample block *b_n_* and a SNP block *b_m_*, and each block *y* in phenotype matrix *Y* be indexed by a sample block *b_n_*.

#### Algorithm 1 Map each block index (*b_n_, b_m_*) to the corresponding genotype matrix block *g* and phenotype matrix block *y*, then emit the products *g^T^g* and *g^T^y*

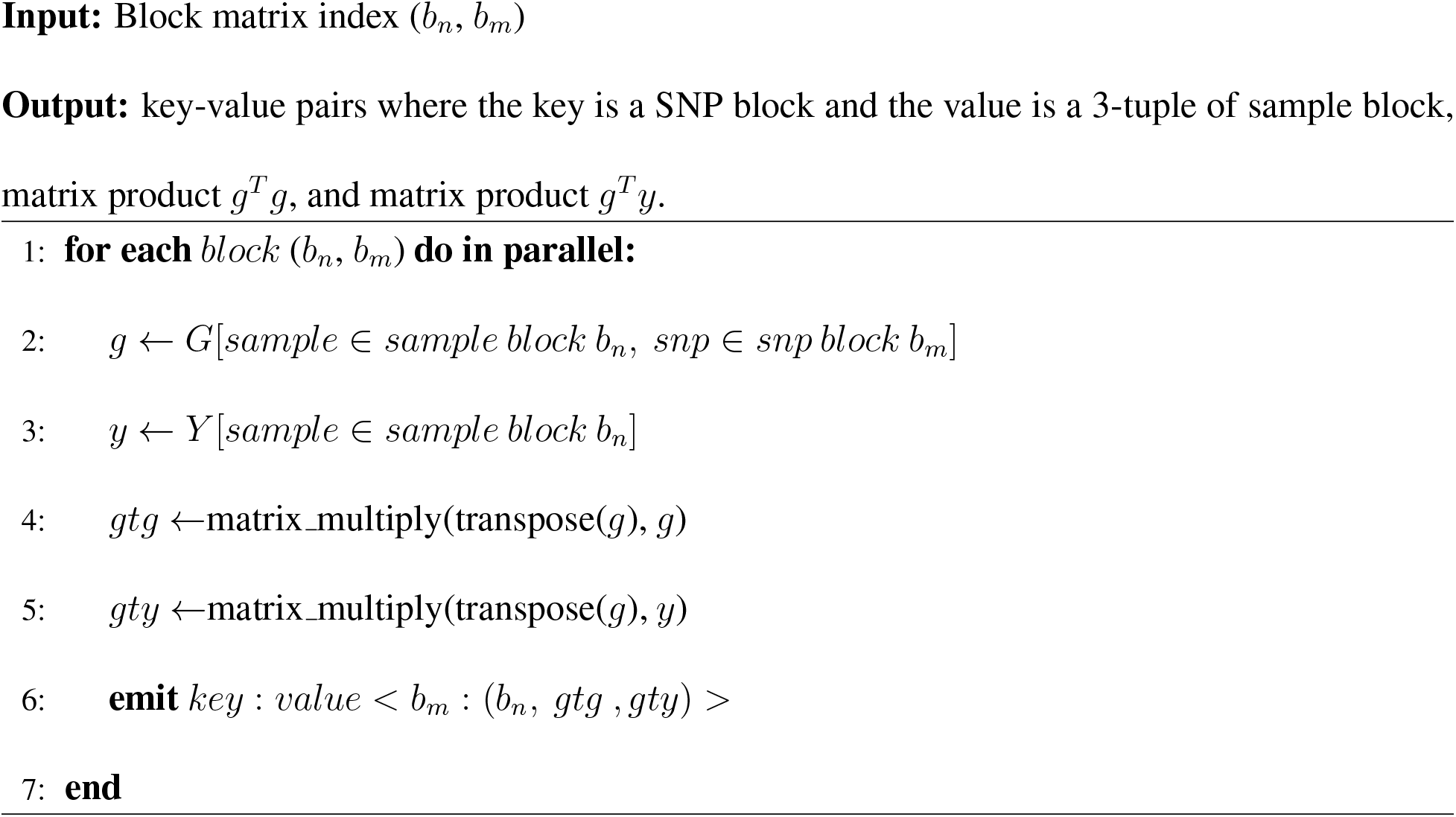

#### Algorithm 2 For each key *b_m_* and value (*b_n_,g^T^g,g^T^y*) pair emitted from Algorithm 1, generate new keys corresponding to block matrix indices (*b_n_, b_m_*) and new values representing the element-wise sums over *g^T^g* and *g^T^y* where sample group *b_n_* has been left out

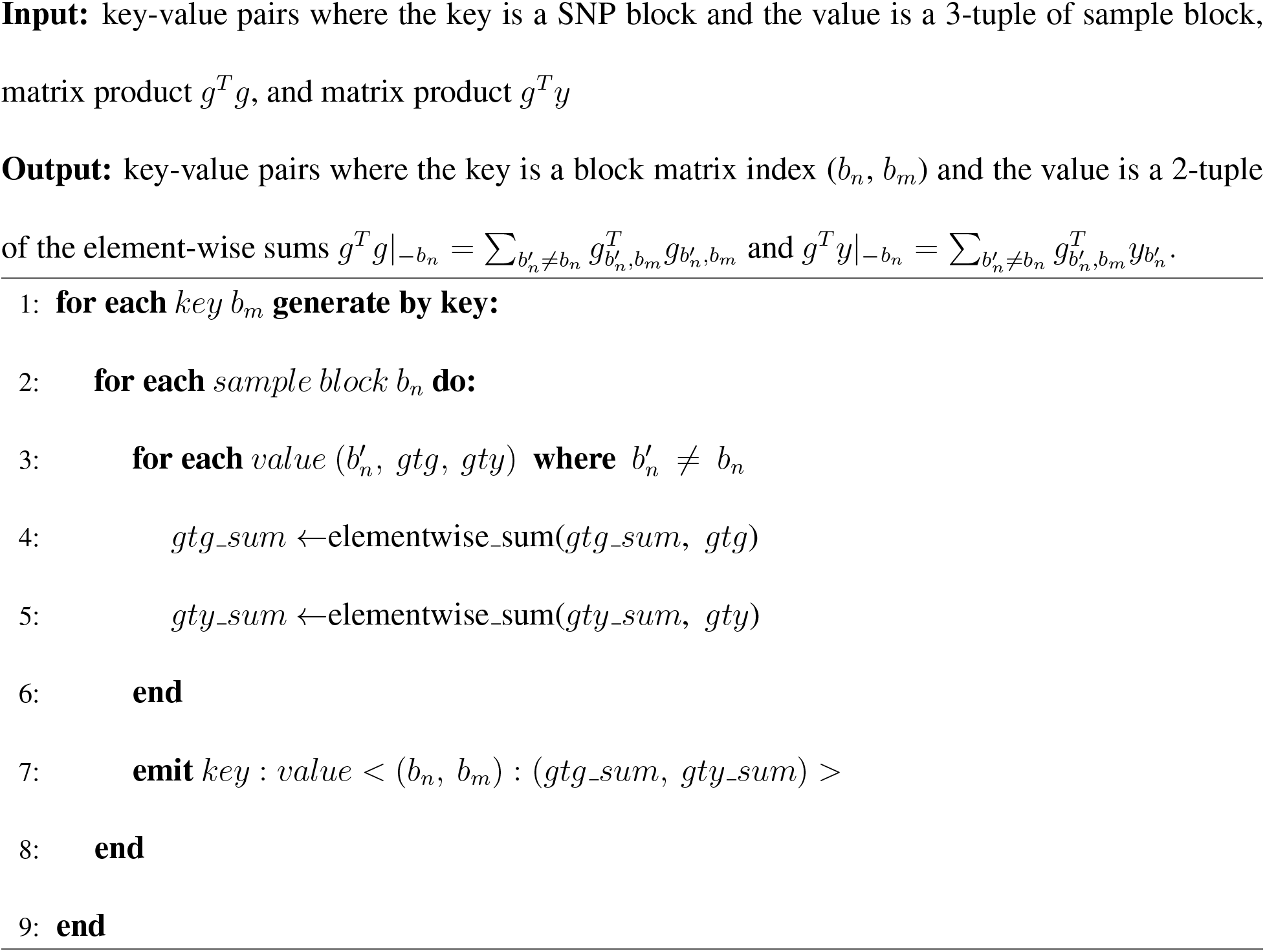

#### Algorithm 3 For each key (*b_n_, b_m_*) emitted from Algorithm 2, map value (*g^T^g*|_−*b_n_*_, *g^T^y*|_−*b_n_*_) to a new value with sample block *b_n_* and ridge predictor matrix block w corresponding to block (*b_n_, b_m_*) with key equal to SNP block *b_m_*

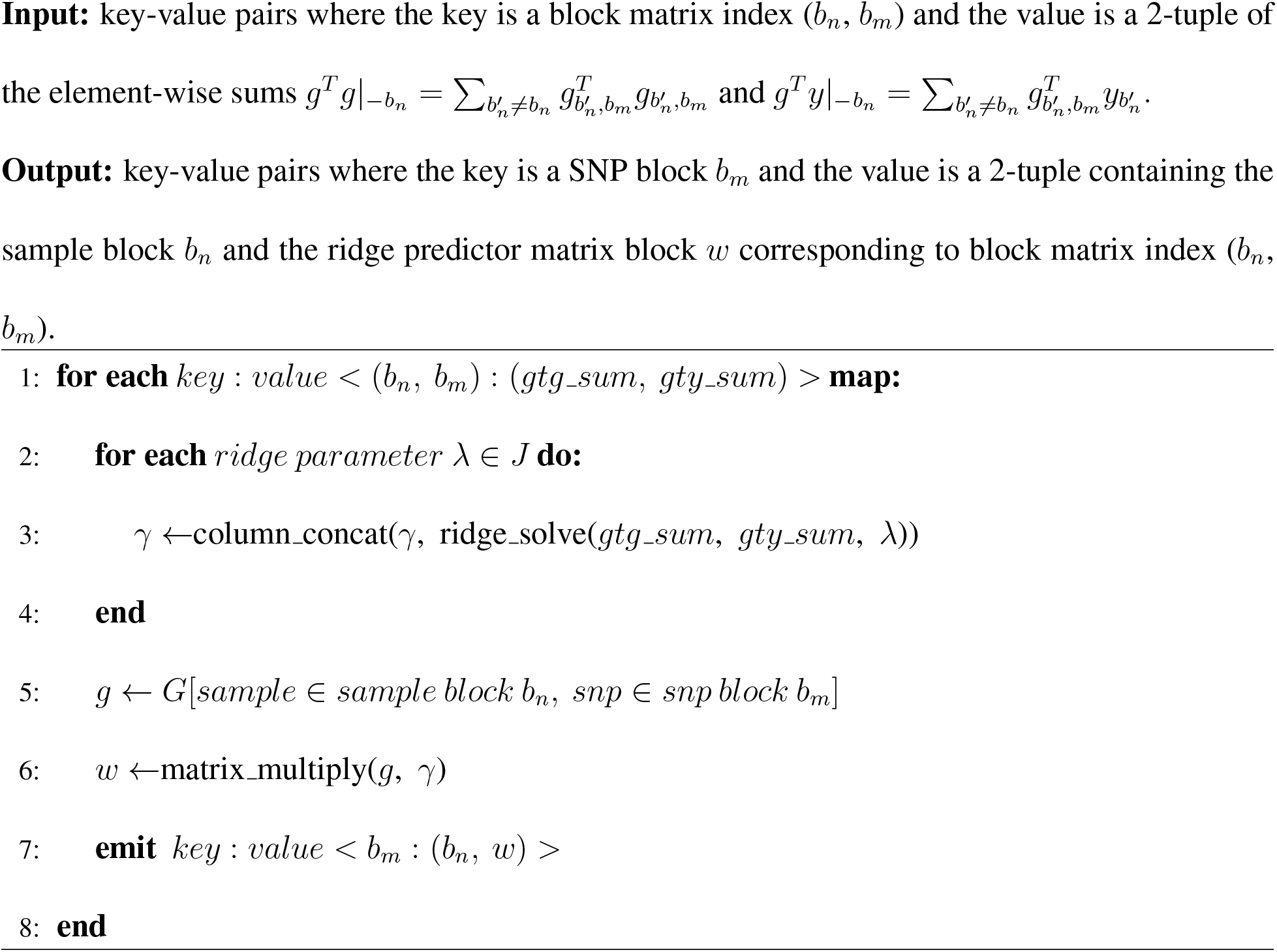

#### Algorithm 4 For each key *b_m_* emitted from Algorithm 3, reduce values (*b_n_, w*) by row-wise concatenation of matrices w

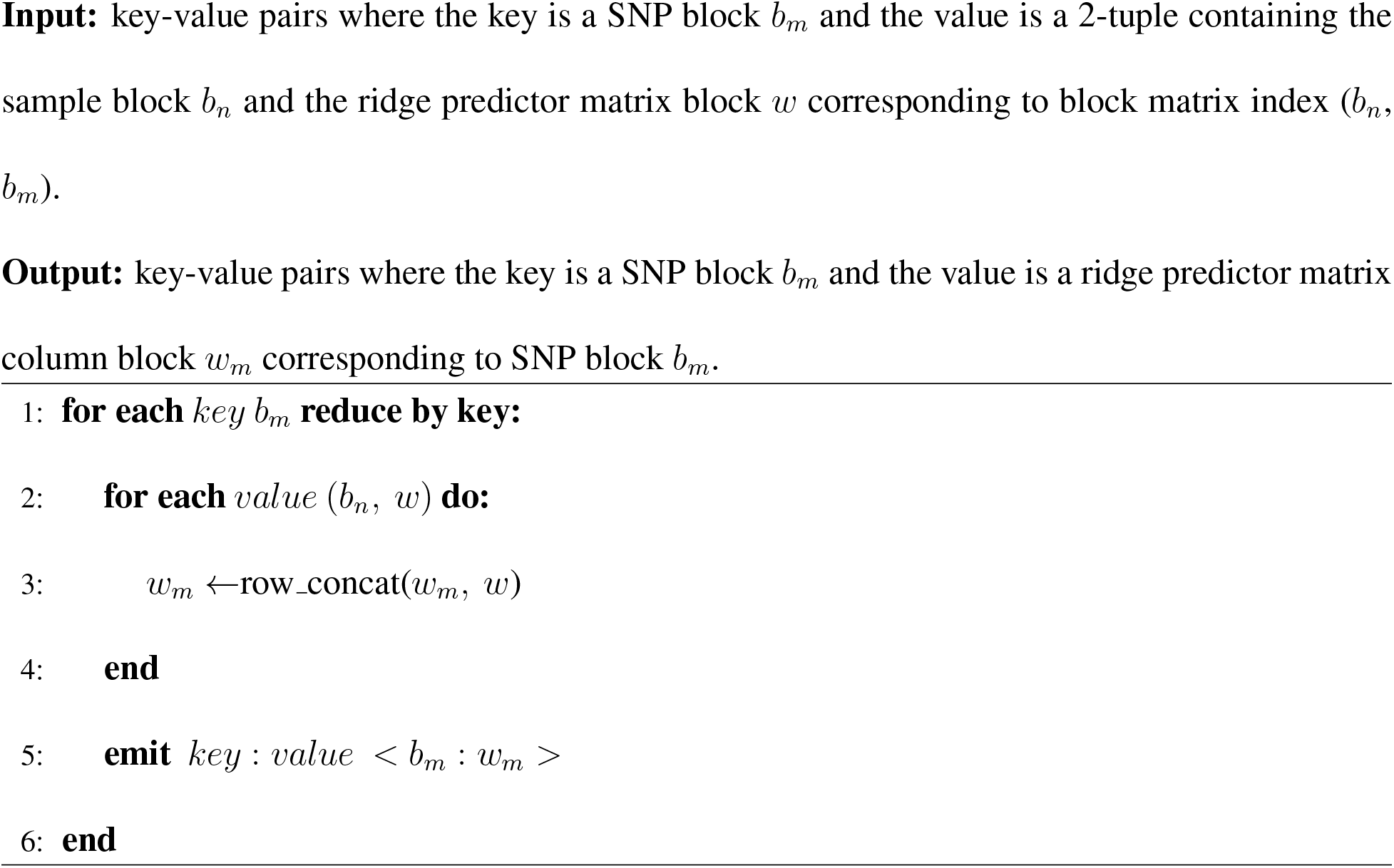

#### Algorithm 5 Reduce values *w_m_* emitted from Algorithm 4 by column-wise concatenation and return ridge predictor matrix *W*

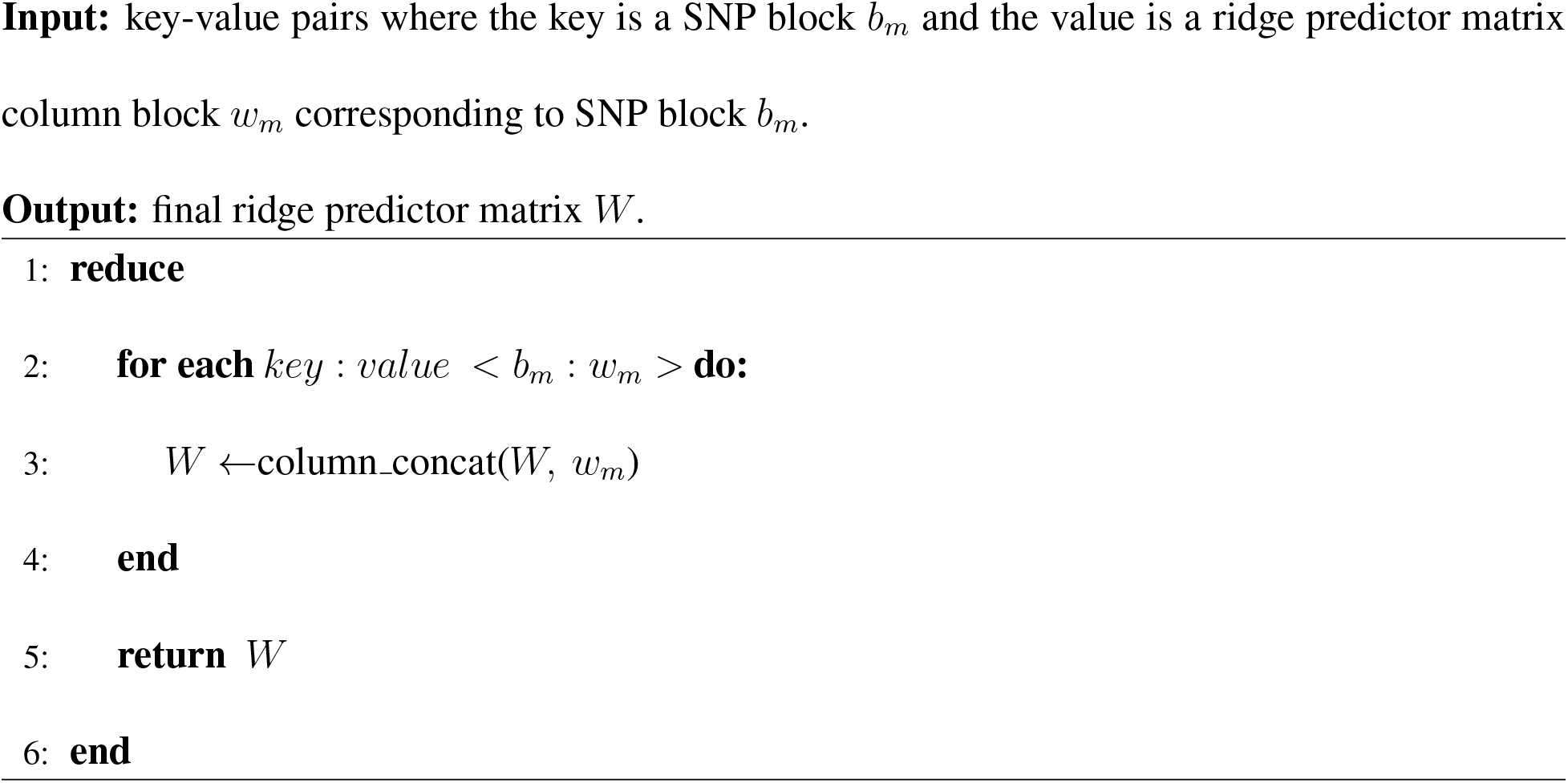

### Inter-chromosomal LD in the UK Biobank

To identify how prevalent inter-chromosome LD is in the UK Biobank array genotypes, we used LDstore^42^ (version 2.0beta) to compute pairwise Pearson correlation coefficients for a set of 343,784 genotyped SNPs with MAF ≥ 0.05, Hardy-Weinberg equilibrium test not exceeding 10^−12^ significance and missingness rate below 5%, in a random sample of 5,000 unrelated white British individuals. This resulted in over 59 billion SNP-pairs for which we computed correlation, among which about 56 billion corresponded to SNP-pairs on different chromosomes. Using Bonferroni correction with a significance level of 5% resulted in a significance threshold of 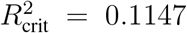 to identify significant SNP-pairs on different chromosomes. **Supplementary Figure S21** displays the 3,697 SNP pairs that were found and that spanned a total of 12 chromosomes. The full list of 3,697 SNP-pairs is available in the **Supplementary Table 10** and the 50 SNP-pairs with the highest correlation coefficients are listed in **Supplementary Table S8**. For many of the SNP-pairs involved, they corresponded to known gene/pseudogene pairs (see **Supplementary Table S9**). For the remaining SNP pairs, it could be that the gene/pseudogene pairs involved have yet to be annotated.

### Supplementary Tables

**Table S1:**
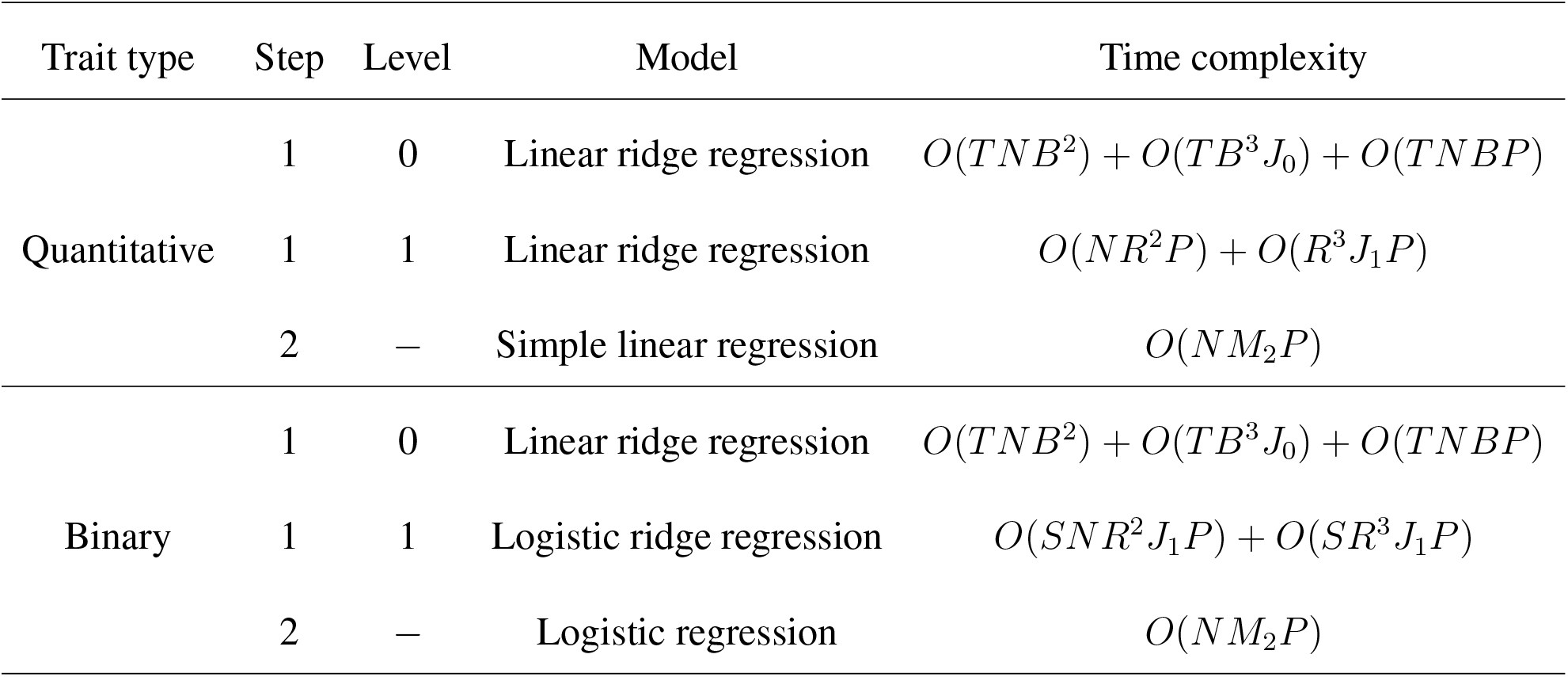
Time complexity of the REGENIE method. Step 1 level 0 is the within-blocks models and Step 1 level 1 is the across-blocks models. Step 2 is the association testing step.

*M*_1_, number of variants used in step 1;
*M*_2_, number of variants tested in step 2;
N, sample size (assumed to be the same for all traits);
*P*, number of phenotypes;
*B*, number of SNPs in a genotype block;
*J*_0_, number of ridge parameters at level 0;
*J*_1_, number of ridge parameters at level 1;
*T* = *M*_1_/*B*, total number of genotype blocks;
*R* = *J*_0_*T*, number of predictors from level 0 models;
*S*, number of iterations to reach convergence in logistic ridge regression.

**Table S2:** Phenotype information for the 50 quantitative traits analyzed using UK Biobank data of 407,746 white British samples. The percentage missing reported is for the full 407K sample.

| Trait# | Phenotype | N | %missing |
| --- | --- | --- | --- |
| 1 | Townsend Deprivation Index At Recruitment | 407,271 | 0.1 |
| 2 | Body Mass Index Bmi | 407,609 | 0.0 |
| 3 | Body Fat Percentage | 401,772 | 1.5 |
| 4 | White Blood Cell Leukocyte Count | 396,621 | 2.7 |
| 5 | Red Blood Cell Erythrocyte Count | 396,625 | 2.7 |
| 6 | Haemoglobin Concentration | 396,624 | 2.7 |
| 7 | Mean Corpuscular Volume | 396,624 | 2.7 |
| 8 | Platelet Count | 396,621 | 2.7 |
| 9 | Mean Platelet Thrombocyte Volume | 396,616 | 2.7 |
| 10 | Lymphocyte Count | 395,949 | 2.9 |
| 11 | Monocyte Count | 395,949 | 2.9 |
| 12 | Neutrophil Count | 395,949 | 2.9 |
| 13 | Eosinophil Count | 395,949 | 2.9 |
| 14 | Basophil Count | 395,949 | 2.9 |
| 15 | Creatinine Enzymatic In Urine | 396,837 | 2.7 |
| 16 | Potassium In Urine | 395,995 | 2.9 |
| 17 | Sodium In Urine | 396,020 | 2.9 |
| 18 | Albumin | 357,968 | 12.2 |
| 19 | Alkaline Phosphatase | 389,883 | 4.4 |
| 20 | Alanine Aminotransferase | 389,733 | 4.4 |
| 21 | Apolipoprotein A | 355,859 | 12.7 |
| 22 | Apolipoprotein B | 388,022 | 4.8 |
| 23 | Peak Expiratory Flow Pef | 374,599 | 8.1 |
| 24 | Aspartate Aminotransferase | 388,490 | 4.7 |
| 25 | Direct Bilirubin | 332,739 | 18.4 |
| 26 | Urea | 389,608 | 4.5 |
| 27 | Calcium | 357,831 | 12.2 |
| 28 | Cholesterol | 389,864 | 4.4 |
| 29 | Creatinine | 389,678 | 4.4 |
| 30 | C Reactive Protein | 389,057 | 4.6 |
| 31 | Cystatin C | 389,834 | 4.4 |
| 32 | Gamma Glutamyltransferase | 389,672 | 4.4 |
| 33 | Glucose | 357,580 | 12.3 |
| 34 | Glycated Haemoglobin Hba1c | 389,889 | 4.4 |
| 35 | Hdl Cholesterol | 357,810 | 12.2 |
| 36 | Igf 1 | 387,834 | 4.9 |
| 37 | Ldl Direct | 389,189 | 4.5 |
| 38 | Phosphate | 357,302 | 12.4 |
| 39 | Shbg | 354,620 | 13.0 |
| 40 | Total Bilirubin | 388,303 | 4.8 |
| 41 | Testosterone | 353,805 | 13.2 |
| 42 | Triglycerides | 389,562 | 4.5 |
| 43 | Urate | 389,404 | 4.5 |
| 44 | Vitamin D | 373,045 | 8.5 |
| 45 | Diastolic Blood Pressure Automated Reading | 385,801 | 5.4 |
| 46 | Systolic Blood Pressure Automated Reading | 385,798 | 5.4 |
| 47 | Hand Grip Strength Left | 406,552 | 0.3 |
| 48 | Waist Circumference | 407,661 | 0.0 |
| 49 | Hip Circumference | 407,662 | 0.0 |
| 50 | Bone Mineral Density | 365,403 | 10.4 |

**Table S3:** Comparison of runtimes for REGENIE model fitting step when analyzing 50 quantitative traits with UK Biobank data of 407,746 individuals. REGENIE was used in multi-trait mode analyzing all traits together either storing block predictions in memory (REGENIE MT) or writing them to disk (REGENIE MT-WD). 469,336 LD-pruned SNPs were used as model SNPs when fitting the null model. All runs were done on the same computing environment (64 virtual CPU cores of a 2.5 GHz Intel Xeon Platinum 8175M processor, 512GB of memory and 2.4TB solid-state disk) and REGENIE was run using up to 16 threads.

| Method | Benchmark |  |  |
| --- | --- | --- | --- |
|  | CPU Time (hours) | Elapsed time (hours) | Memory usage (GB) |
| REGENIE MT | 62 | 9.4 | 390 |
| REGENIE MT-WD | 79 | 11 | 13 |

**Table S4:** Comparison of rare variant effect size estimates from different methods. Results shown for thyroid cancer (case-control ratio 1:660) using UK Biobank data of 407,746 white British samples. Estimates for the top 10 imputed SNPs with the most discrepancy between REGENIE and SAIGE are displayed. The estimates are compared between REGENIE using approximate Firth correction, SAIGE, which uses SPA correction, and PLINK using Firth correction. For PLINK, only covariates are adjusted for when testing each variant for association (i.e. no polygenic effect predictions are included in the model).

| SNP | MAC | Estimated effect $\hat{\beta}$ (log OR) | | | SE($\hat{\beta}$) | | |
| --- | --- | --- | --- | --- | --- | --- | --- |
|  |  | REGENIE | SAIGE | PLINK | REGENIE | SAIGE | PLINK |
| 4:82827568:G:A | 5.7 | -8.1 | -468.1 | -7.4 | 1.5 | 125.4 | 1.4 |
| 5:131807265:A:G | 5.4 | -8.3 | -708.8 | -8.7 | 2.2 | 184.2 | 2.3 |
| 11:55936336:C:A | 19.0 | -6.5 | -381.4 | -6.8 | 1.4 | 106.0 | 1.4 |
| 11:18446202:A:T | 5.3 | -10.0 | -1039.3 | -13.4 | 3.7 | 269.7 | 3.6 |
| 1:228148490:C:T | 5.1 | -11.1 | -985.6 | -13.7 | 3.8 | 259.5 | 3.8 |
| 3:12156938:A:G | 5.7 | -10.9 | -424.9 | -10.6 | 2.3 | 136.2 | 2.3 |
| 1:161998091:T:A | 6.3 | -9.4 | -370.7 | -9.2 | 2.3 | 131.2 | 2.4 |
| 10:125833896:A:C | 7.5 | -8.3 | -287.3 | -7.5 | 2.0 | 110.5 | 2.1 |
| 6:152168123:C:T | 8.2 | -4.7 | -233.4 | -6.4 | 1.3 | 68.6 | 1.3 |
| 5:38406872:G:T | 6.5 | -8.4 | -454.1 | -8.2 | 2.2 | 121.1 | 1.9 |

**Table S5:** Comparison of run time of exact and approximate Firth logistic regression. Runtimes for REGENIE association testing step when analyzing 4 binary traits with UK Biobank data of 407,746 individuals. REGENIE was run using either the exact Firth correction (REGENIE-FIRTH Exact) or using an approximation of that test which adjusts for covariates through an offset term in the model (REGENIE-FIRTH Approx). 516,196 imputed variants were tested for association with each of the four binary traits. All runs were done on the same computing environment (16 virtual CPU cores of a 2.5 GHz Intel Xeon Platinum 8175M processor and 64GB of memory).

| Method | Benchmark |  |  |
| --- | --- | --- | --- |
|  | CPU Time (hours) | Elapsed time (hours) | Memory usage (GB) |
| REGENIE-FIRTH Exact | 5232 | 420 | 4.8 |
| REGENIE-FIRTH Approx | 87 | 7.5 | 2.8 |

**Table S6:** Phenotype information for the 50 binary traits analyzed using UK Biobank data of 407,746 white British samples. The percentage missing reported is for the full 407K sample.

| Trait# | Phenotype | N | CC ratio | %missing |
| --- | --- | --- | --- | --- |
| 1 | Mood Swings | 407,746 | 1:1 | 0.0 |
| 2 | Miserableness | 407,746 | 1:1 | 0.0 |
| 3 | Irritability | 407,746 | 1:3 | 0.0 |
| 4 | Guilty Feelings | 407,746 | 1:2 | 0.0 |
| 5 | Risk Taking | 407,746 | 1:3 | 0.0 |
| 6 | Seen Doctor For Nerves Anxiety Tension Or Depression | 407,746 | 1:2 | 0.0 |
| 7 | Hearing Difficulty Problems | 407,746 | 1:3 | 0.0 |
| 8 | Dd Vascular Heart Problems 4 High Blood Pressure | 407,746 | 1:3 | 0.0 |
| 9 | Pain Types Experienced In Last Month 1 Headache | 407,746 | 1:4 | 0.0 |
| 10 | Allergicrhinitis | 407,746 | 1:3 | 0.0 |
| 11 | High Cholesterol | 407,746 | 1:7 | 0.0 |
| 12 | Asthma | 401,847 | 1:7 | 1.4 |
| 13 | Seen Psych Nerves Anxiety Tension Or Depression | 407,746 | 1:8 | 0.0 |
| 14 | Chest Pain Or Discomfort | 407,746 | 1:5 | 0.0 |
| 15 | Fractured Bones In Last 5 Years | 407,746 | 1:9 | 0.0 |
| 16 | Reason For Glasses Contact Lenses 1 For Short Sightedness | 407,746 | 1:8 | 0.0 |
| 17 | Pain Types Experienced In Last Month 6 Hip Pain | 407,746 | 1:7 | 0.0 |
| 18 | Medication For Cholesterol Blood Pressure Or Diabetes | 407,746 | 1:8 | 0.0 |
| 19 | Any Parental History Of Alzheimers | 407,089 | 1:7 | 0.2 |
| 20 | Any Parental History Of Lung Cancer | 407,521 | 1:8 | 0.1 |
| 21 | Depression | 407,746 | 1:17 | 0.0 |
| 22 | Osteoarthritis | 407,746 | 1:11 | 0.0 |
| 23 | Angina | 397,554 | 1:17 | 2.5 |
| 24 | Cholelithiasis Gall Stones | 404,405 | 1:21 | 0.8 |
| 25 | Urinary Tract Infection Kidney Infection | 397,867 | 1:18 | 2.4 |
| 26 | Type 2 Diabetes | 406,831 | 1:20 | 0.2 |
| 27 | Hypothyroidism Myxoedema | 405,357 | 1:16 | 0.6 |
| 28 | Back Problem | 402,528 | 1:19 | 1.3 |
| 29 | Inflammatory Bowel Disease | 404,781 | 1:24 | 0.7 |
| 30 | Eye Problems Disorders 4 Cataract | 407,746 | 1:24 | 0.0 |
| 31 | Pulmonary Embolism DVT | 407,746 | 1:115 | 0.0 |
| 32 | Epilepsy | 407,746 | 1:117 | 0.0 |
| 33 | Retinal Detachment | 403,643 | 1:100 | 1.0 |
| 34 | Muscle Soft Tissue Problem | 407,746 | 1:117 | 0.0 |
| 35 | Allergy Hypersensitivity Anaphylaxis | 407,498 | 1:102 | 0.1 |
| 36 | Bronchitis | 407,746 | 1:116 | 0.0 |
| 37 | Atrial Fibrillation | 407,746 | 1:111 | 0.0 |
| 38 | Cystitis | 381,591 | 1:112 | 6.4 |
| 39 | Retinitis Pigmentosa | 403,833 | 1:115 | 1.0 |
| 40 | Macular Degeneration | 403,837 | 1:106 | 1.0 |
| 41 | Bells Palsy Facial Nerve Palsy | 407,746 | 1:1072 | 0.0 |
| 42 | Spinal Cord Disorder | 407,746 | 1:1092 | 0.0 |
| 43 | Sjogrens Syndrome Sicca Syndrome | 407,746 | 1:1095 | 0.0 |
| 44 | Clotting Disorder Excessive Bleeding | 407,746 | 1:1122 | 0.0 |
| 45 | Thyroid Goitre | 407,746 | 1:1069 | 0.0 |
| 46 | Insomnia | 407,746 | 1:1078 | 0.0 |
| 47 | Psoriasis | 407,746 | 1:991 | 0.0 |
| 48 | Type 1 Diabetes | 407,746 | 1:1034 | 0.0 |
| 49 | Acute Infective Polyneuritis Guillain Barre Syndrome | 406,298 | 1:1088 | 0.4 |
| 50 | Cellulitis | 407,746 | 1:1101 | 0.0 |

**Table S7:** Carbon footprint of REGENIE-FIRTH, REGENIE-SPA, and SAIGE when analyzing 50 binary traits with UK Biobank data. Carbon emissions represent the amount of CO_2_ that would have the same impact on global warming as that due to the mix of gases generated from running the software. Carbon sequestration measures how long it would take for a mature tree to absorb the CO_2_ generated from running the software. Car distance measures the travel distance of a passenger car that would emit the same amount of CO_2_ as that generated from running the software. The last column represents the equivalent round-trip car distance between New York (NY) and other states in the US. The calculations were performed based on a cloud computing environment with 16 virtual CPU cores of a 2.1GHz AMD EPYC 7571 processor with a thermal design power of 15W per core. The usage factor was estimated as (CPU hours / Elapsed hours)/16, with the runtimes obtained from **Table 2** where we used the results based on leave-one-out crossvalidation (LOOCV) for REGENIE.

| Method | Carbon emissions<br>(kg CO<sub>2</sub>e) | Carbon sequestration<br>(tree-years) | Car distance<br>(km) | From NY<br>to |
| --- | --- | --- | --- | --- |
| REGENIE-FIRTH | 349 | 15 | 1390 | Virginia |
| REGENIE-SPA | 220 | 10 | 876 | New Hampshire |
| SAIGE | 2,780 | 122 | 11,073 | California |

**Table S8:**
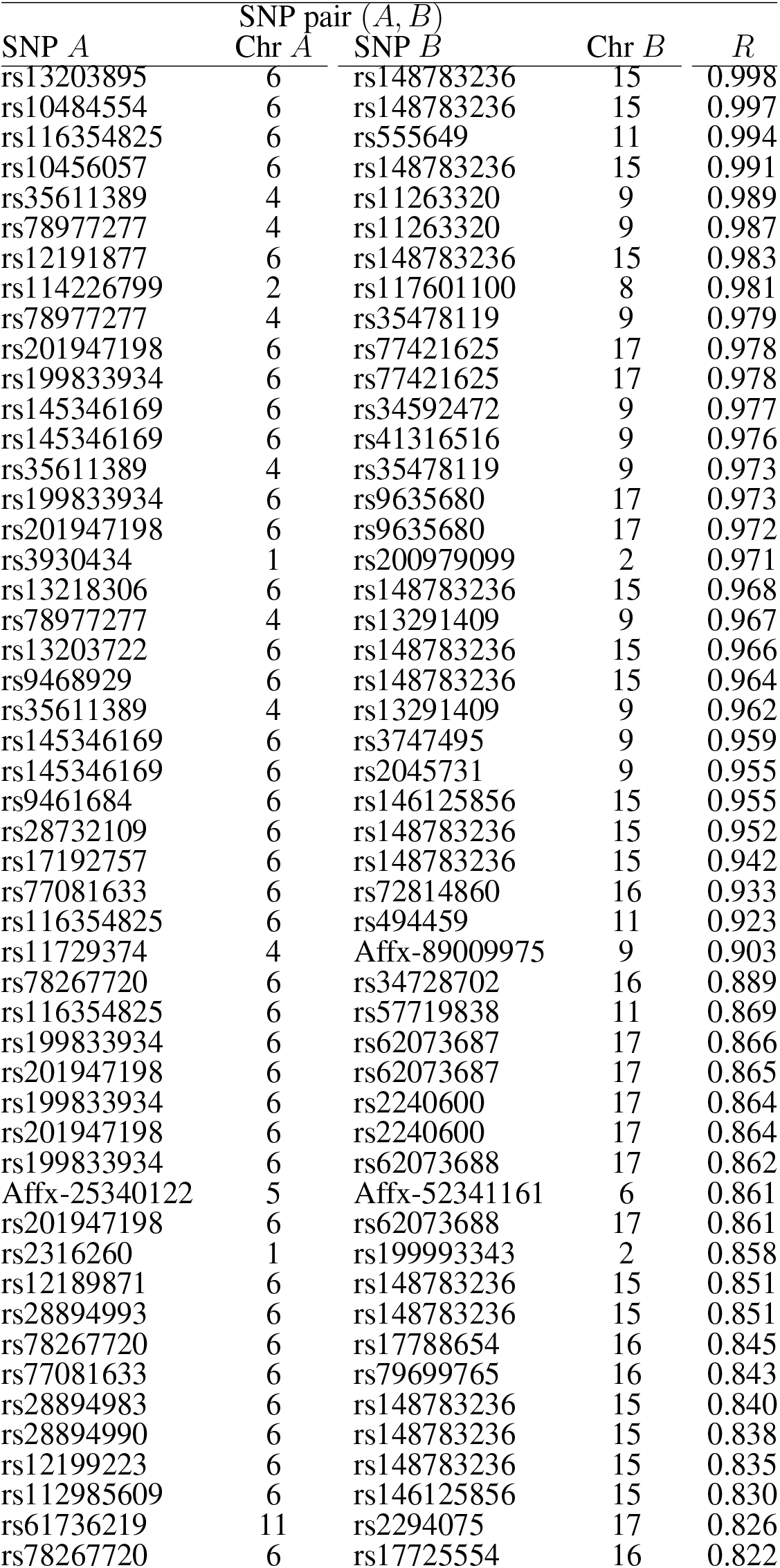
List of 50 inter-chromosome LD (ICLD) SNP pairs from UK Biobank array data with the highest amount of correlation. The Pearson correlation coefficient R is obtained from a sample of 5,000 unrelated white British participants (see **Supplementary Methods**). The full list of 3,697 SNP-pairs is available in **Supplementary Materials**.

**Table S9:** List of gene/pseudogene pairs detected in the inter-chromosome LD (ICLD) analysis of UK Biobank array data.

| Gene (chromosome) | Pseudogene (chromosome) |
| --- | --- |
| *HS6ST1* (2) | HS6ST1P1 (1) |
| *BCLAF1* (6) | *BCLAF1P2*(16) |
| *FOXO3* (6) | *ZNF286B/FOXO3B* (17) |
| *GJA1* (6) | *GJA1P1* (5) |
| *GLDC* (9) | *GLDCP1* (4) |
| *DDX6* (11) | *DDX6P1* (6) |
| *USP8* (15) | *USP8P1* (6) |

### Supplementary Figures

**Figure S1:**
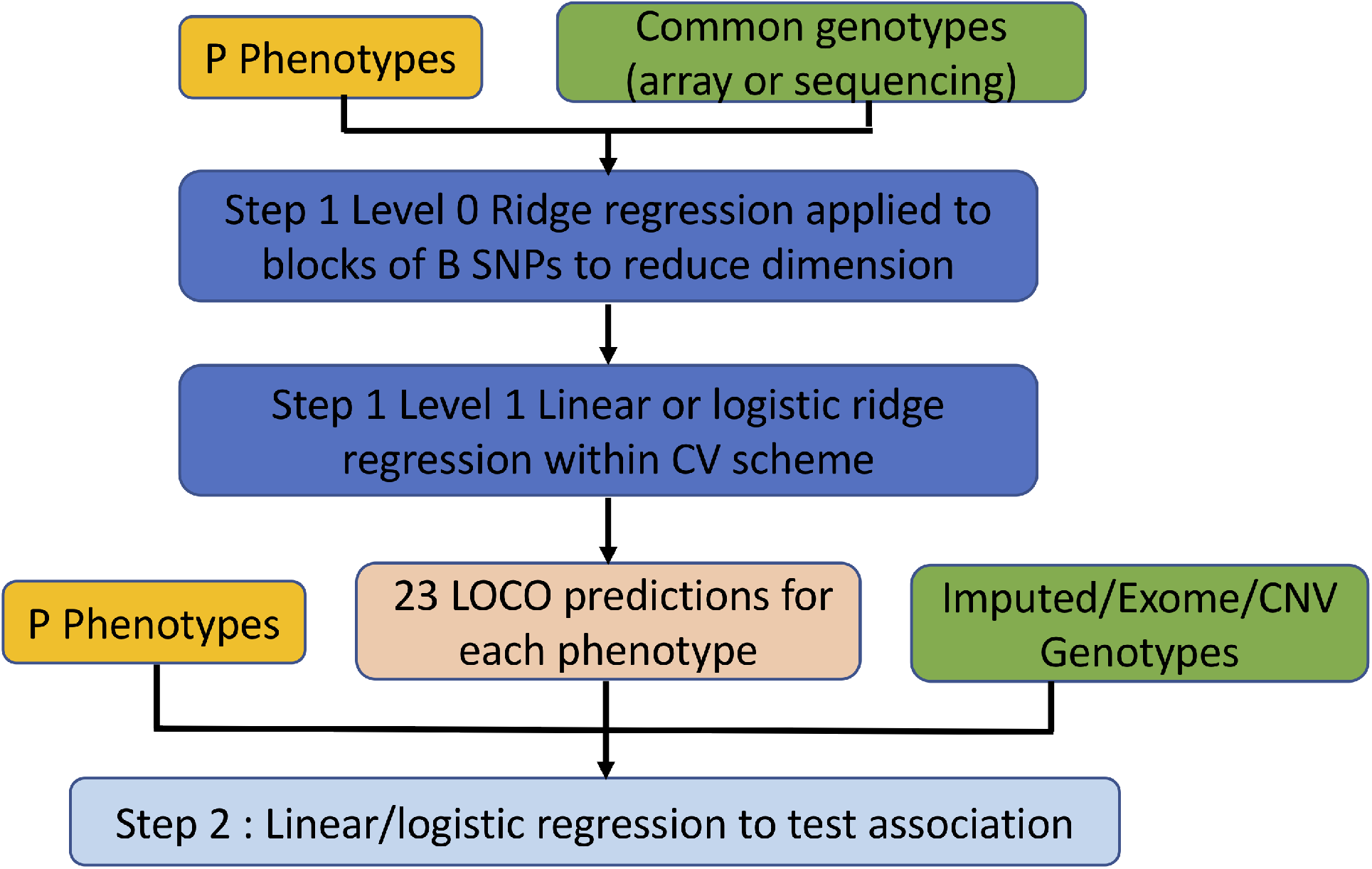
Overview of the REGENIE method. REGENIE consists of two steps: (1) In Step 1, the dimension of the genetic data is reduced using ridge regression applied to blocks of SNPs, and then the resulting predictors are combined using a second round of linear or logistic ridge regression to produce an overall prediction for each trait, split into 23 LOCO predictors. (2) In Step 2, these LOCO predictors are used when testing each phenotype against a set of either imputed, exome or CNV markers.

**Figure S2:**
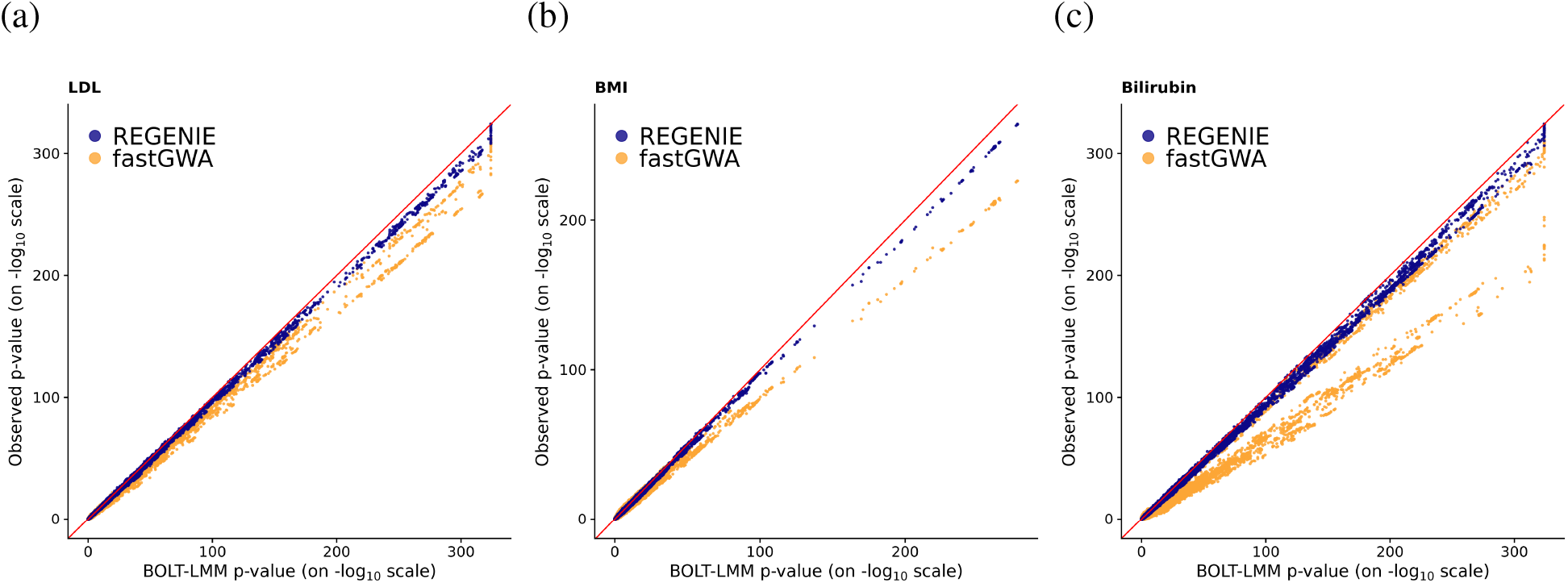
Scatter-plots comparing three LMM methods for three quantitative traits using UK Biobank white British samples. Results from REGENIE, fastGWA and BOLT-LMM are compared for (a) LDL (N = 389, 189), (b) BMI (N = 407, 609) and (c) Bilirubin (N = 388, 303).

**Figure S3:**
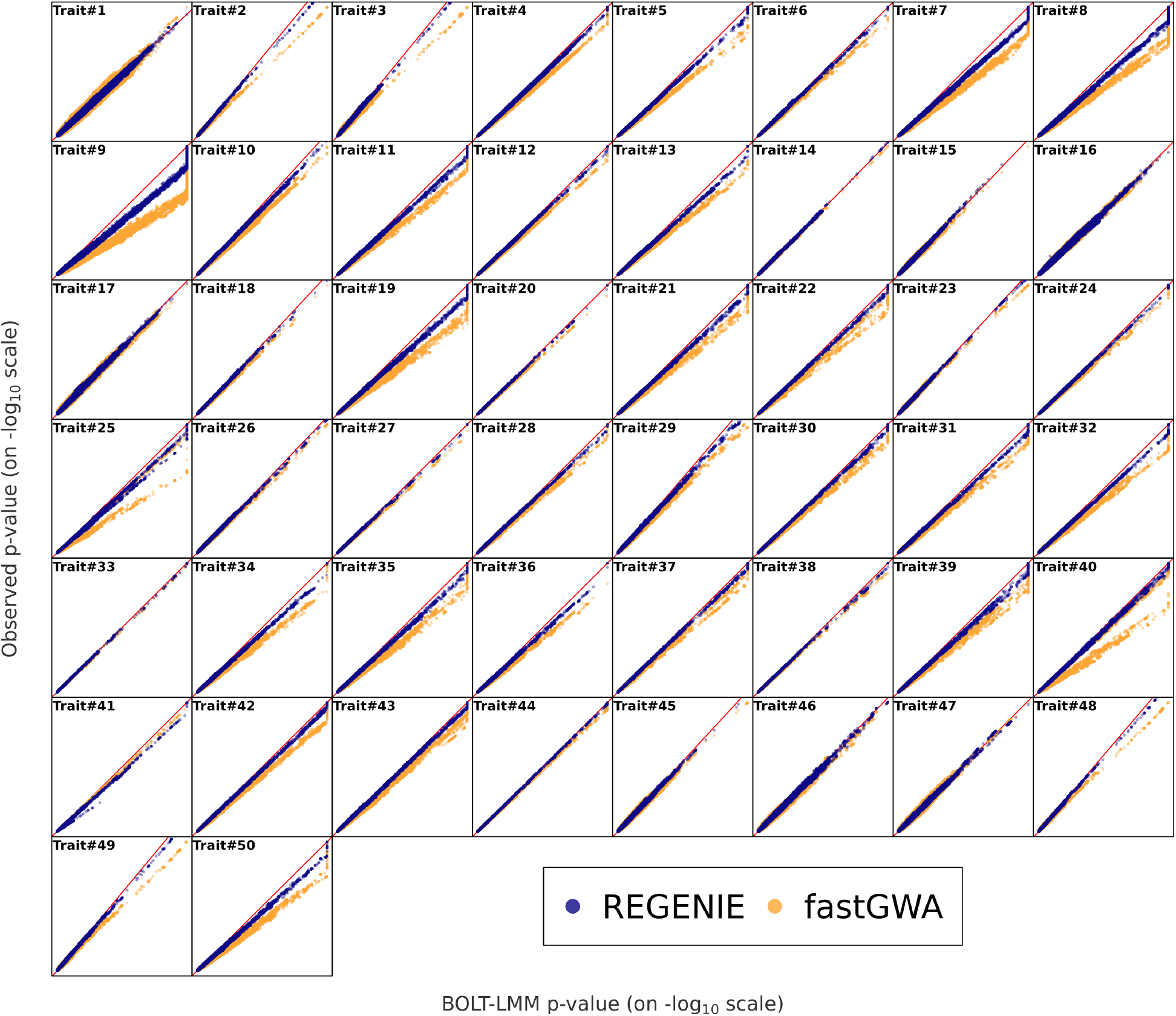
Scatter-plots comparing association results of three LMM methods for 50 quantitative traits using UK Biobank white British samples. 9.8 million imputed SNPs are tested for association with each trait. Information on the traits is available in **Supplementary Table S2**. The scaling of the axes varies over the 50 traits.

**Figure S4:**
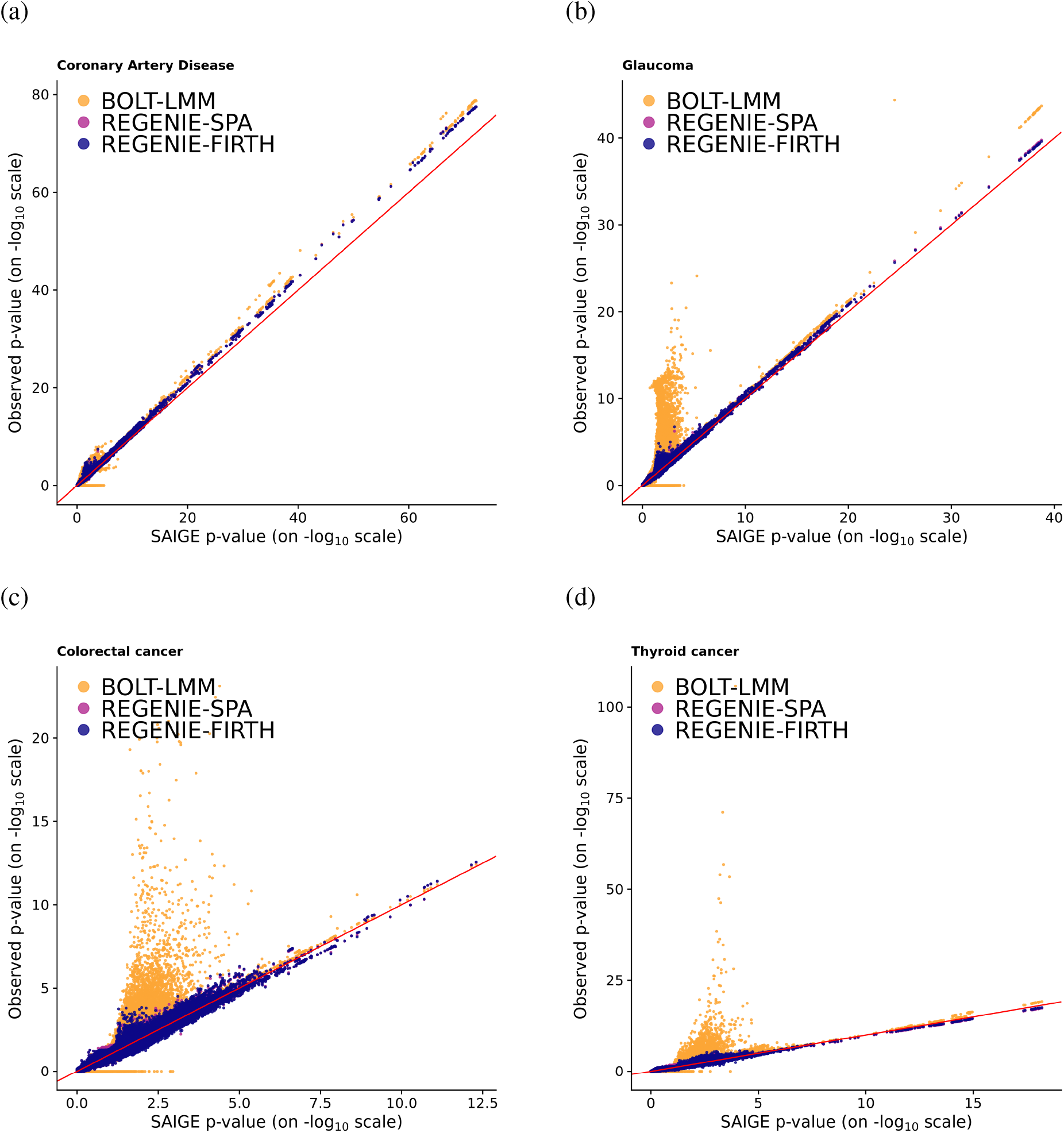
Scatter-plots comparing results from different mixed model methods for 4 binary traits using UK Biobank white British samples. Results from REGENIE using Firth and SPA correction, BOLT-LMM and SAIGE are compared for (a) coronary artery disease (case-control ratio=1:11, N = 352, 063), (b) glaucoma (case-control ratio=1:52, N = 406, 927), (c) colorectal cancer (case-control ratio=1:97, N= 407, 746), and (d) thyroid cancer (case-control ratio=1:660, N= 407, 746). Tests were performed on 11.6 million imputed SNPs.

**Figure S5:**
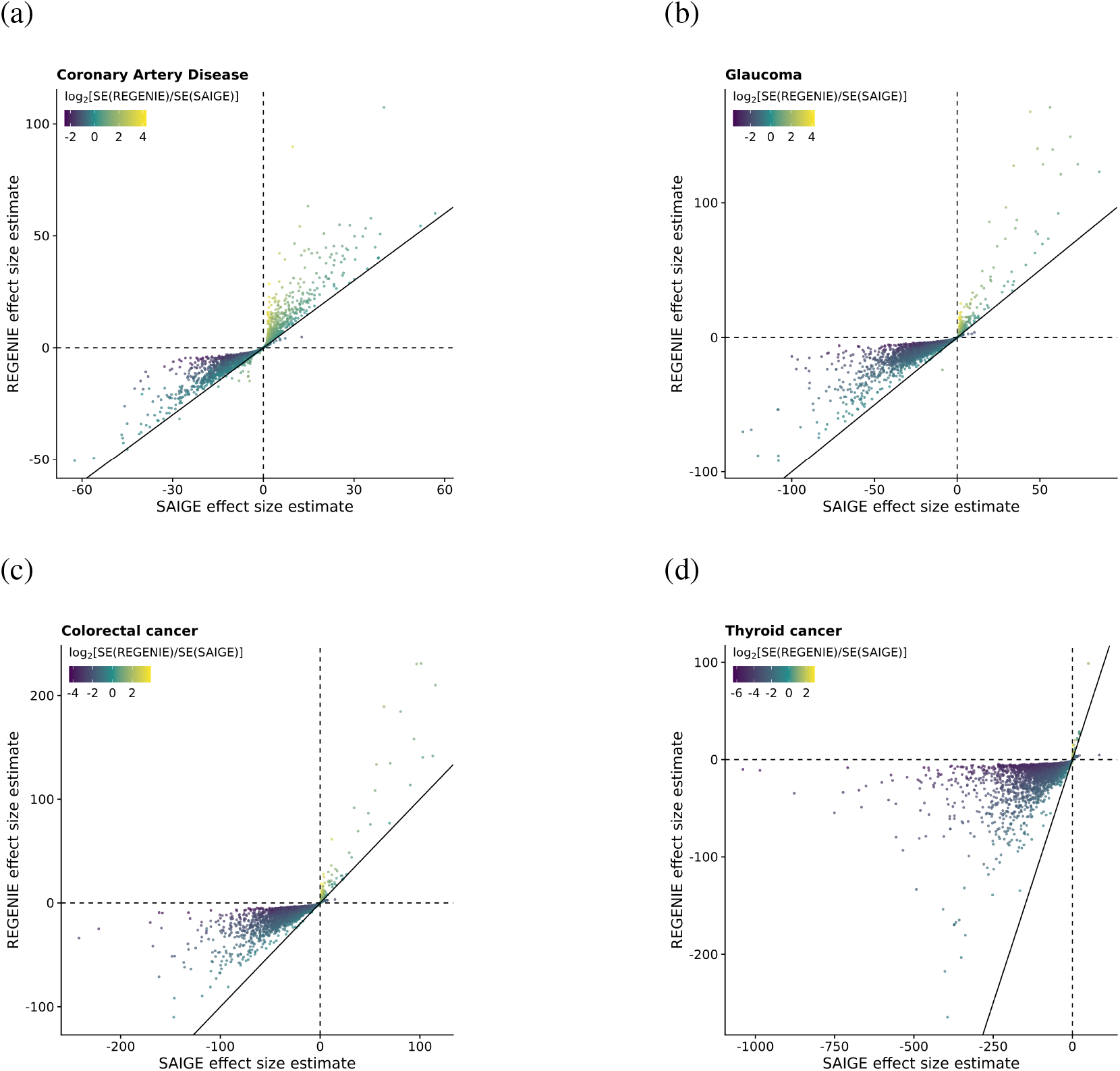
Scatter-plots comparing effect size estimates from two mixed model methods for 4 binary traits using UK Biobank white British samples. Results from REGENIE using Firth correction and SAIGE are compared for (a) coronary artery disease (case-control ratio=1:11, *N*= 352,063), (b) glaucoma (case-control ratio=1:52, N = 406,927), (c) colorectal cancer (case-control ratio=1:97, N= 407,746), and (d) thyroid cancer (case-control ratio=1:660, N= 407,746). Only tests with p-values less than 0.05 in both REGENIE and SAIGE are shown, where 0.05 is the p-value threshold below which Firth/SPA correction is applied. The magnitude of the ratio of the standard errors for the estimated effect sizes from both methods (on log_2_ scale) is represented by the color of the points.

**Figure S6:**
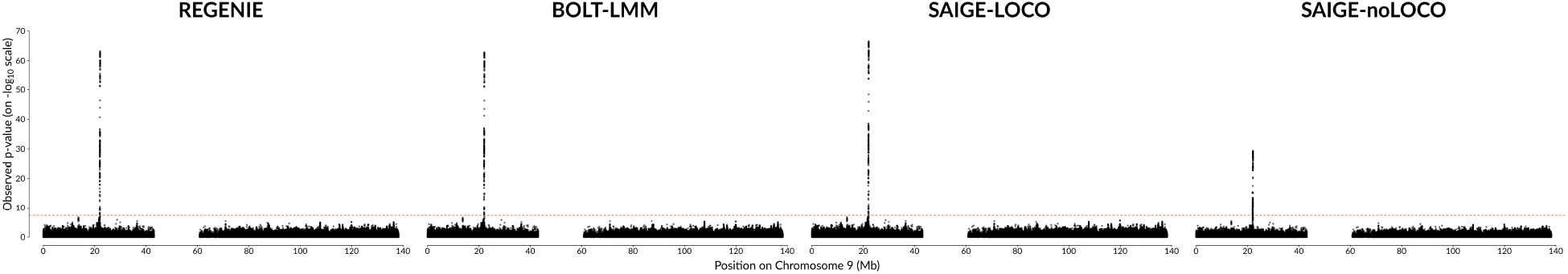
Scatter-plots comparing association results from different mixed model approaches for coronary artery disease using 337,484 unrelated white British participants from UK Biobank. For REGENIE, BOLT-LMM and SAIGE-noLOCO, 329,641 genotyped SNPs from chromosomes 1-22 are included as model SNPs in step 1, and for SAIGE-LOCO all SNPs from chromosome 9 are excluded which results in 314,309 SNPs. In step 2, 482,884 imputed SNPs on chromosome 9 are tested for association. The red dashed horizontal line represents the genomewide significance level of 5 × 10^−8^.

**Figure S7:**
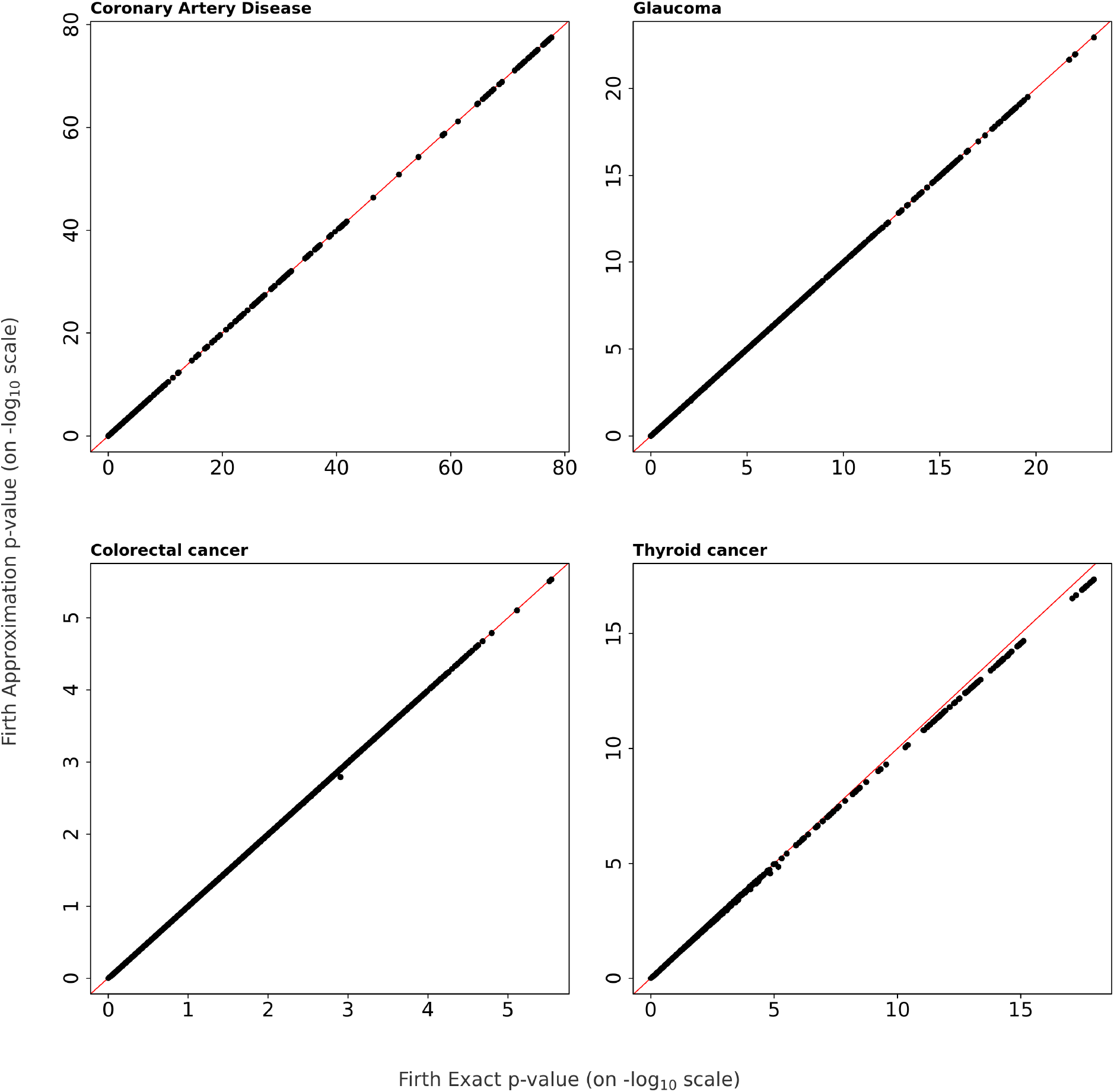
Scatter-plots comparing association results from REGENIE using a Firth-based correction for 4 binary traits using UK Biobank white British samples. Step 2 of REGENIE was run using the Firth correction (Firth Exact) and it was also run using an approximation to the Firth correction (Firth Approximation). 504,441 imputed SNPs on chromosome 9 are tested for association with each of the traits.

**Figure S8:**
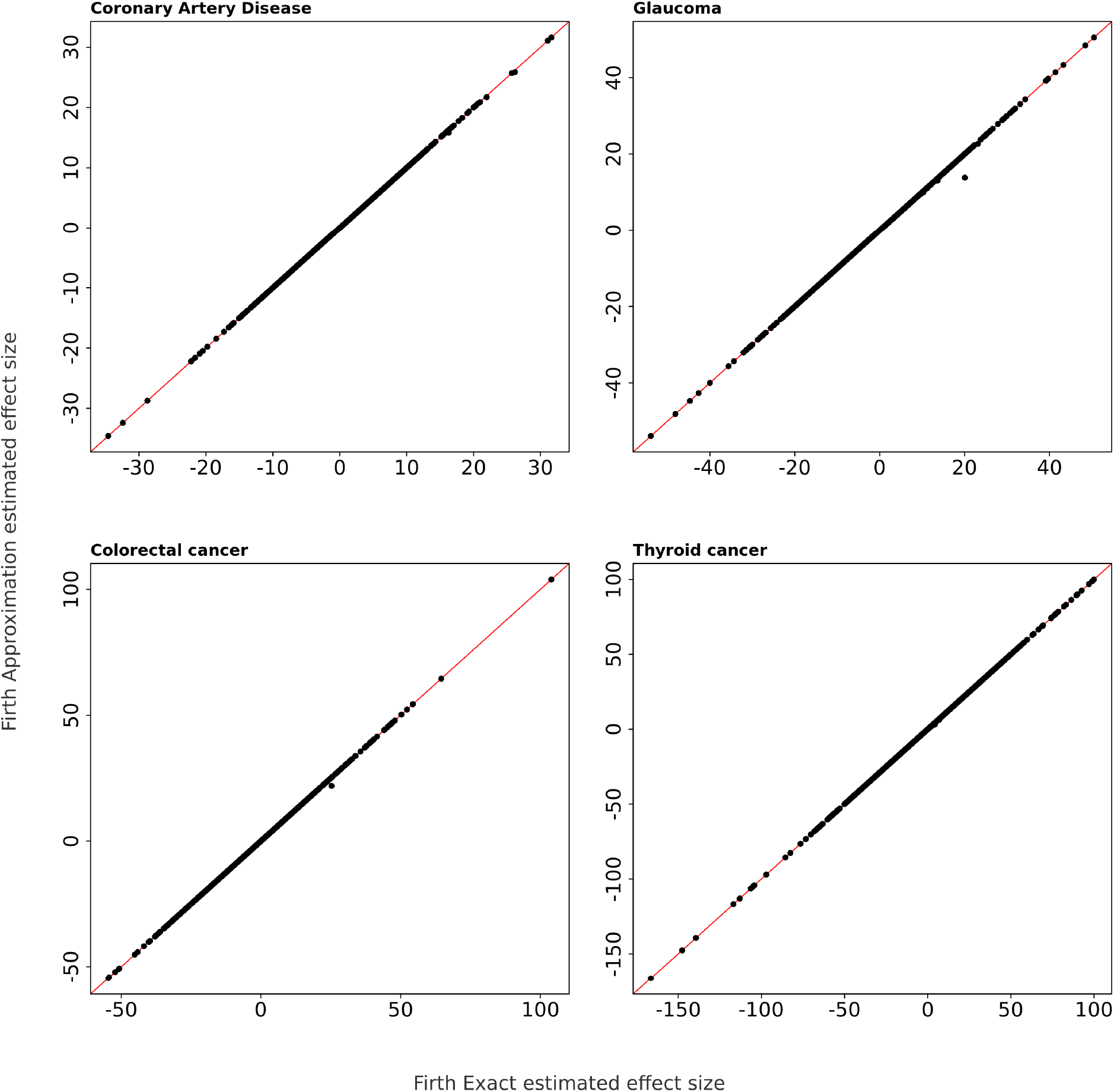
Scatter-plots comparing effect size estimates from REGENIE using a Firth-based correction for 4 binary traits using UK Biobank white British samples. Step 2 of REGENIE was run using the Firth correction (Firth Exact) and it was also run using an approximation to the Firth correction (Firth Approximation). 504,441 imputed SNPs on chromosome 9 are tested for association with each of the traits.

**Figure S9:**
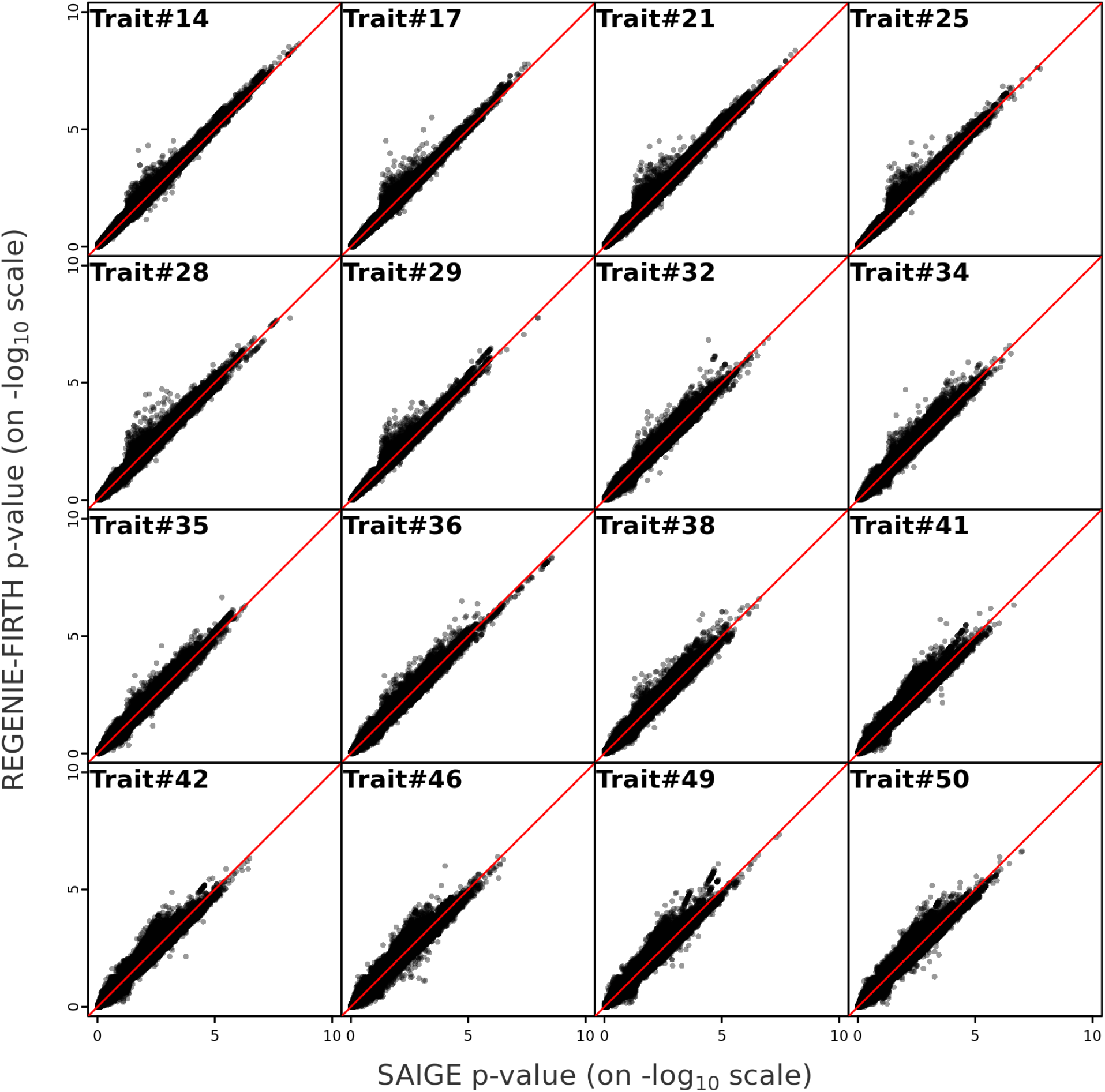
Comparison of REGENIE-FIRTH and SAIGE for 16 of the 50 binary traits using UK Biobank white British samples. 11 million imputed SNPs are tested for association with each trait. Association results are shown for 16 traits with minimum p-value ≥ 10^−10^ for REGENIEFIRTH. Information on the traits is available in **Supplementary Table S6**.

**Figure S10:**
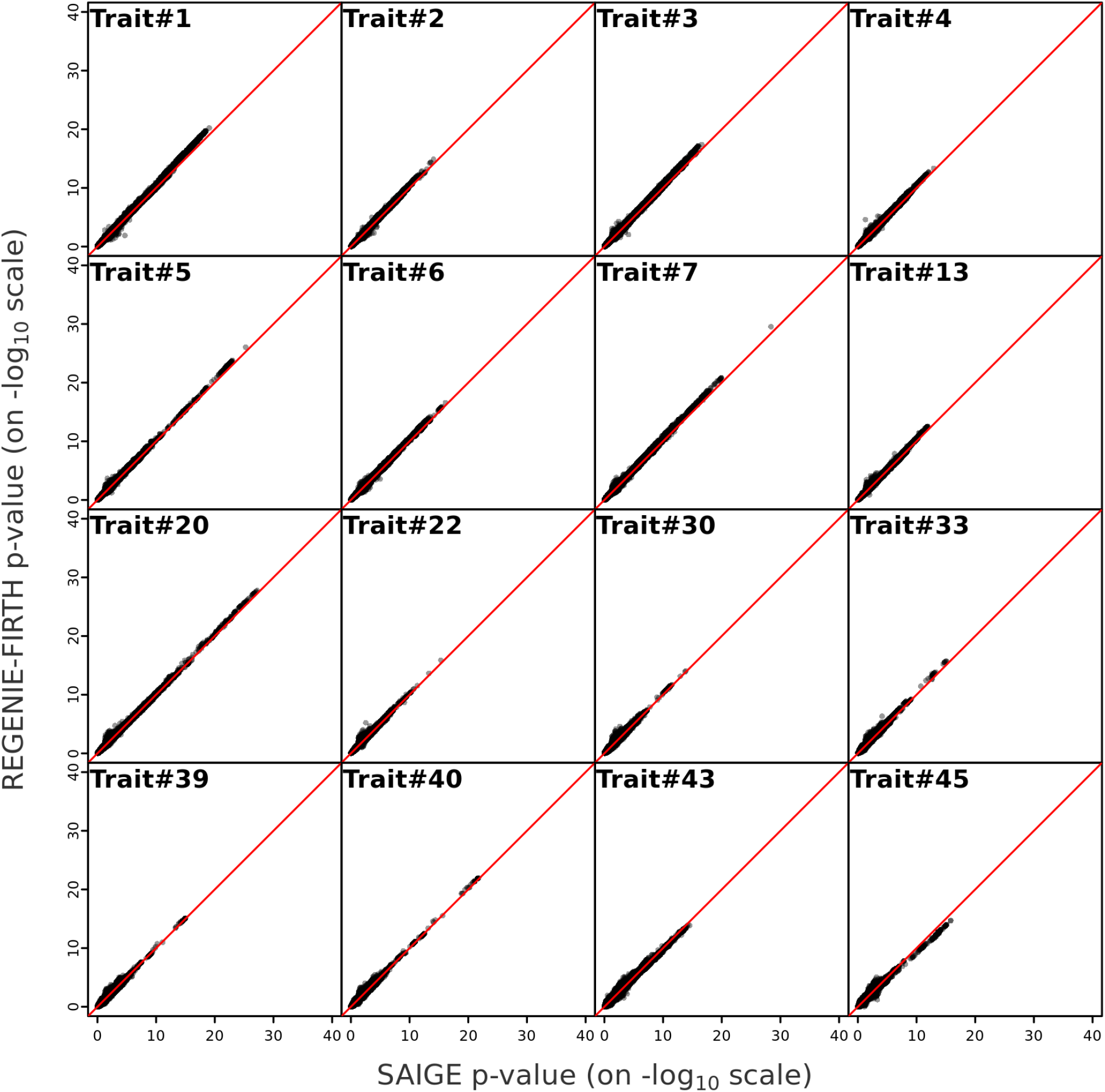
Comparison of REGENIE-FIRTH and SAIGE for 16 of the 50 binary traits using UK Biobank white British samples. 11 million imputed SNPs are tested for association with each trait. Association results are shown for 16 traits with minimum p-value between 10^−10^ and 10^−30^ for REGENIE-FIRTH. Results are omitted for traits which took longer than 4 weeks to fit the null model in SAIGE (DNF = did not finish). Information on the traits is available in **Supplementary Table S6**.

**Figure S11:**
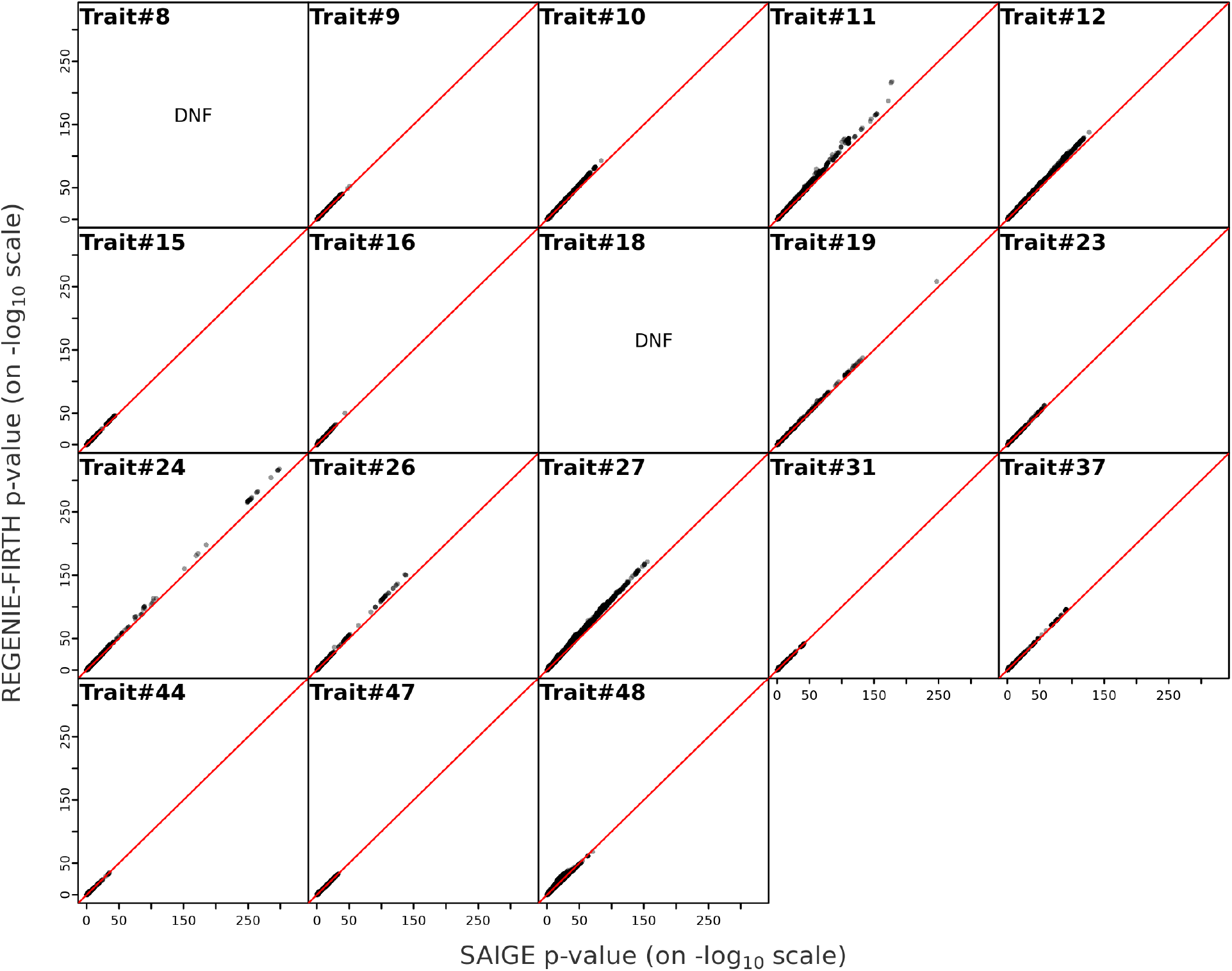
Comparison of REGENIE-FIRTH and SAIGE for 18 of the 50 binary traits using UK Biobank white British samples. 11 million imputed SNPs are tested for association with each trait. Association results are shown for 18 traits with minimum p-value ≤ 10^−30^ for REGENIE-FIRTH. Results are omitted for traits which took longer than 4 weeks to fit the null model in SAIGE (DNF = did not finish). Information on the traits is available in **Supplementary Table S6**.

**Figure S12:**
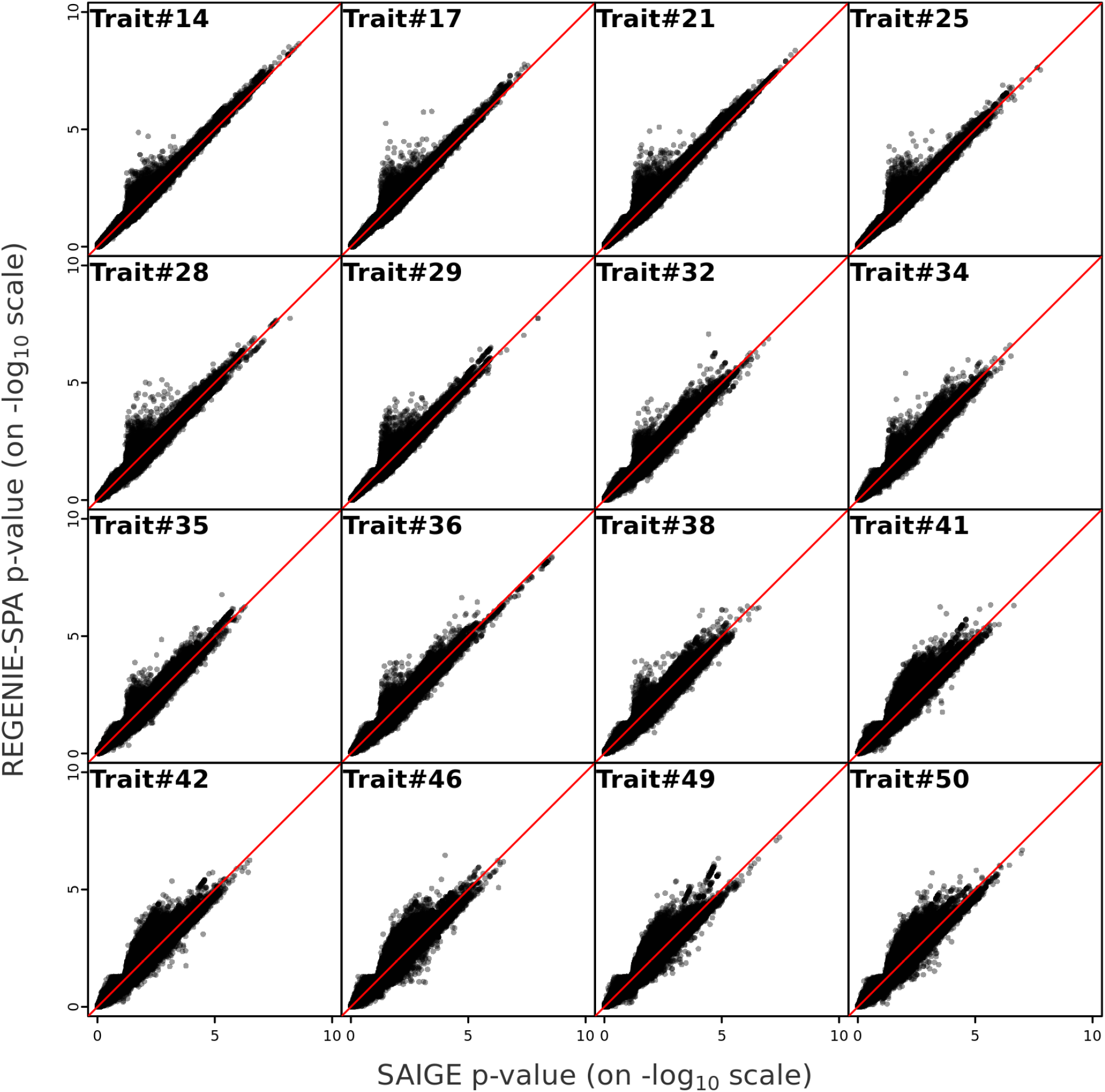
Comparison of REGENIE-SPA and SAIGE for 16 of the 50 binary traits using UK Biobank white British samples. 11 million imputed SNPs are tested for association with each trait. Association results are shown for 16 traits with minimum p-value ≥ 10^−10^ for REGENIESPA. Information on the traits is available in **Supplementary Table S6**.

**Figure S13:**
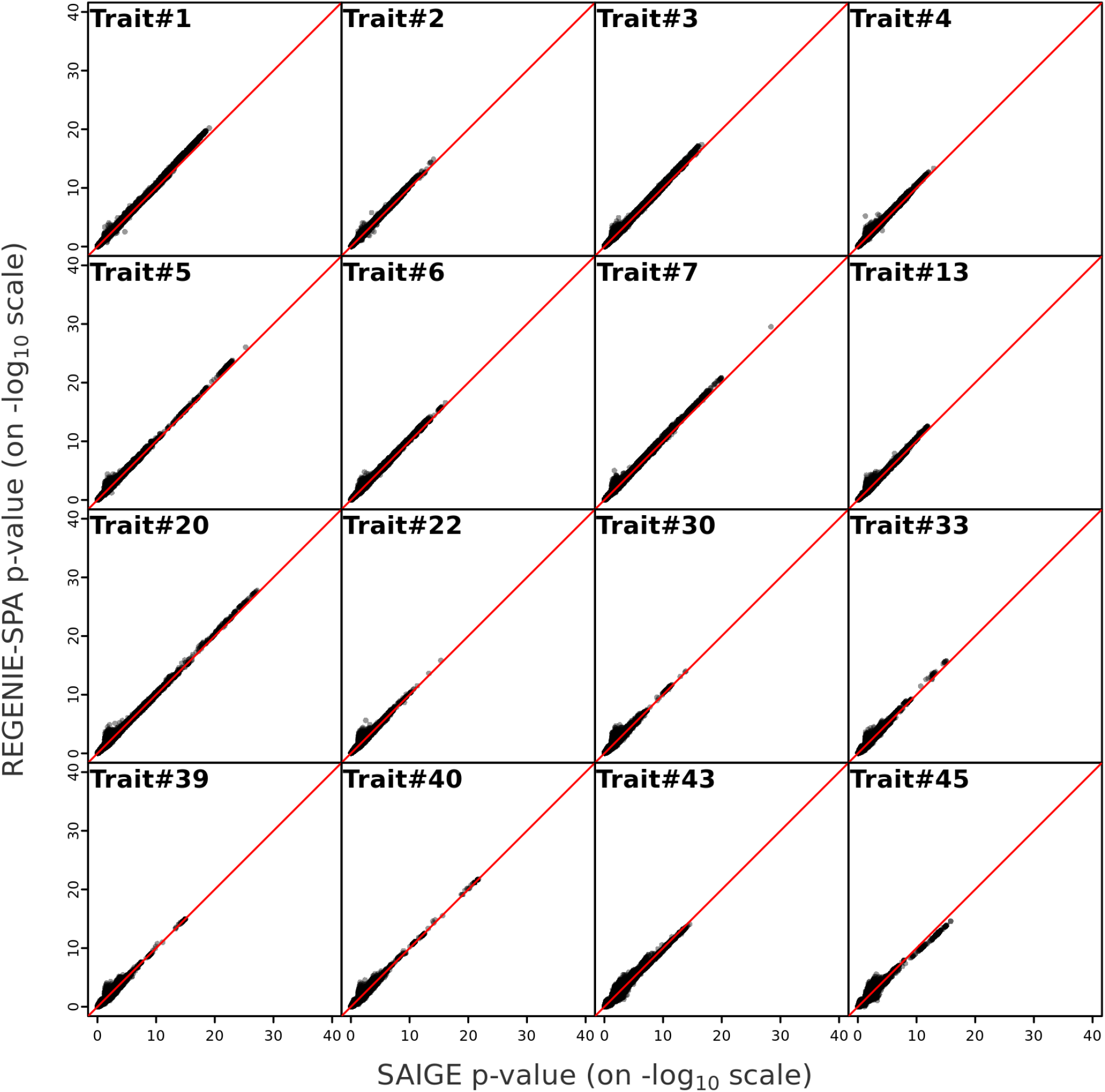
Comparison of REGENIE-SPA and SAIGE for 16 of the 50 binary traits using UK Biobank white British samples. 11 million imputed SNPs are tested for association with each trait. Association results are shown for 16 traits with minimum p-value between 10^−10^ and 10^−30^ for REGENIE-SPA. Results are omitted for traits which took longer than 4 weeks to fit the null model in SAIGE (DNF = did not finish). Information on the traits is available in **Supplementary Table S6**.

**Figure S14:**
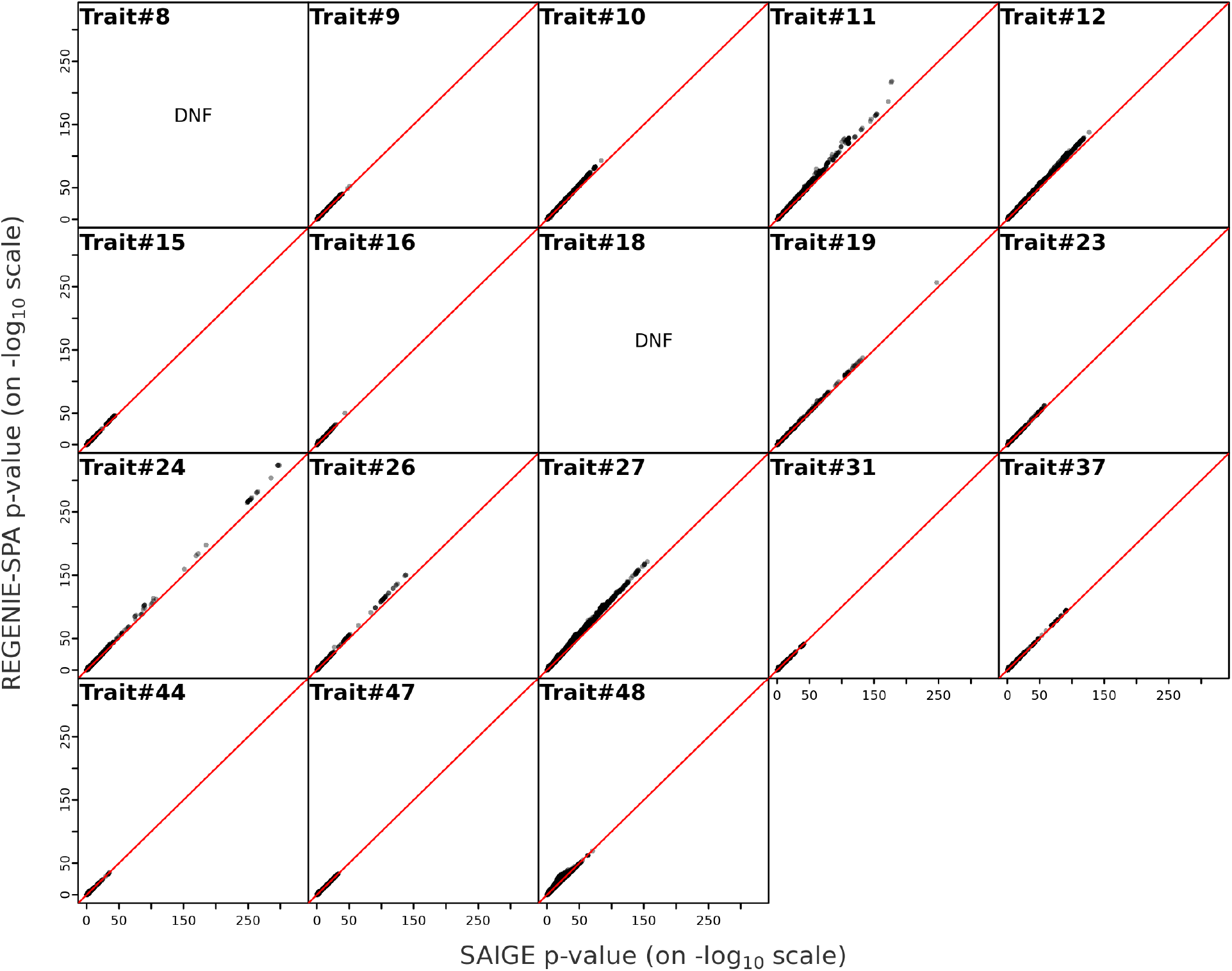
Comparison of REGENIE-SPA and SAIGE for 18 of the 50 binary traits using UK Biobank white British samples. 11 million imputed SNPs are tested for association with each trait. Association results are shown for 18 traits with minimum p-value ≤ 10^−30^ for REGENIESPA. Results are omitted for traits which took longer than 4 weeks to fit the null model in SAIGE (DNF = did not finish). Information on the traits is available in **Supplementary Table S6**.

**Figure S15:**
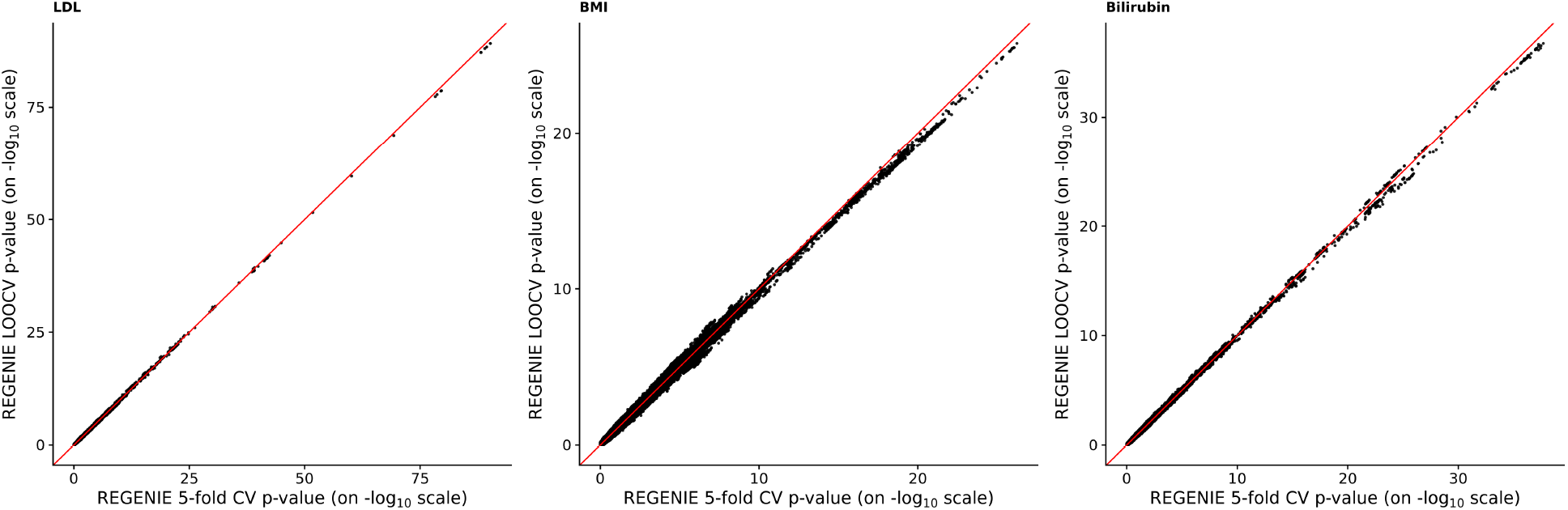
Comparison of REGENIE K-fold and leave-one-out cross validation schemes for quantitative traits. Scatter-plots comparing association results of three quantitative traits using UK Biobank white British samples where null model fitting step of REGENIE is ran with different cross-validation schemes. The null model fitting step 1 of REGENIE is ran using 5-fold cross validation (REGENIE 5-fold CV) and also using leave-one out cross validation (REGENIE LOOCV). 504,441 imputed SNPs on chromosome 9 are tested for association.

**Figure S16:**
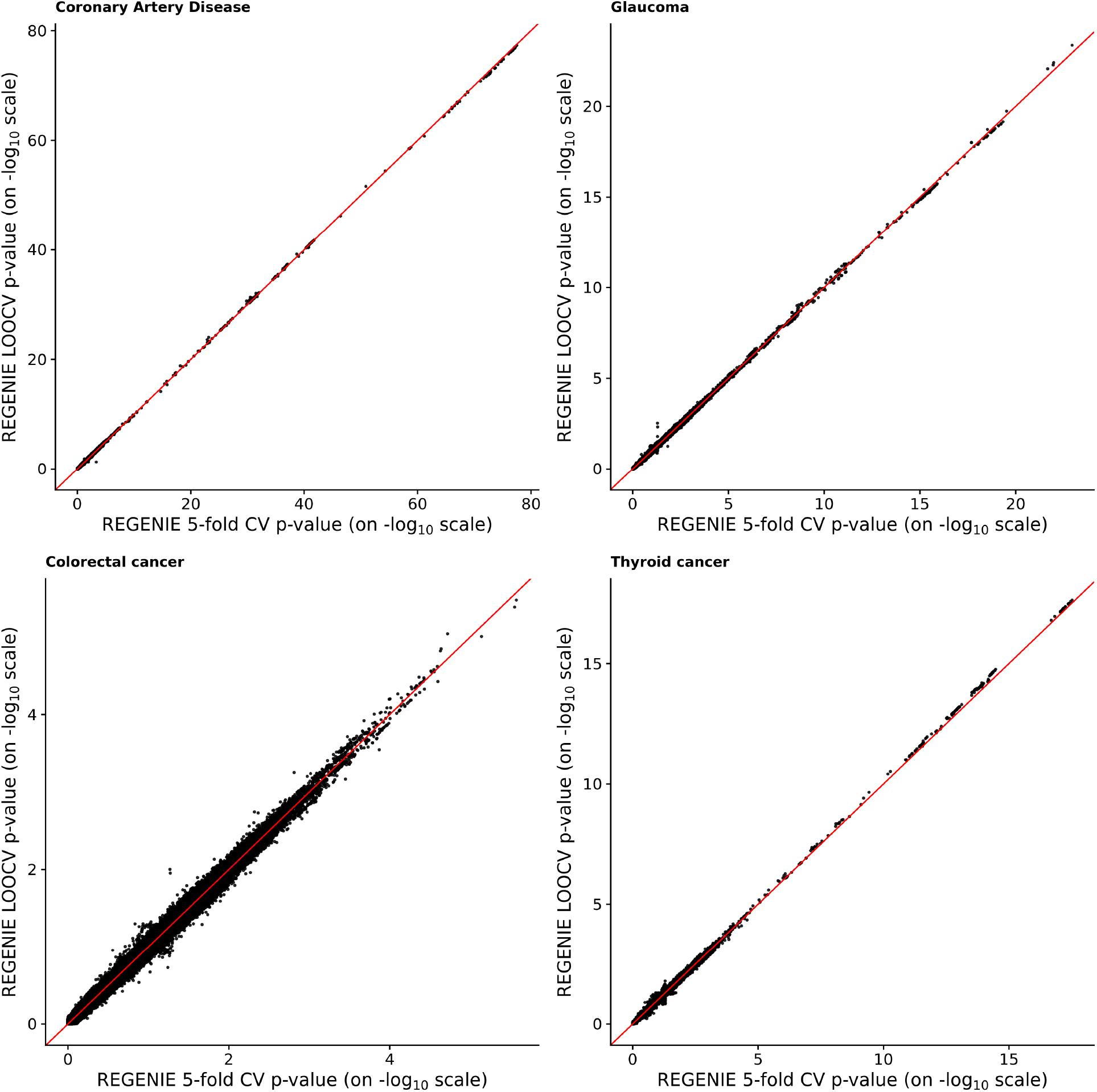
Comparison of REGENIE K-fold and leave-one-out cross validation schemes for binary trait. Scatter-plots compare association results of 4 binary traits using UK Biobank white British samples where null model fitting step of REGENIE was run with different cross-validation schemes. The x-axis is 5-fold cross validation (REGENIE 5-fold CV) and y-axis is leave-one out cross validation (REGENIE LOOCV). 504,441 imputed SNPs on chromosome 9 are tested for association.

**Figure S17:**
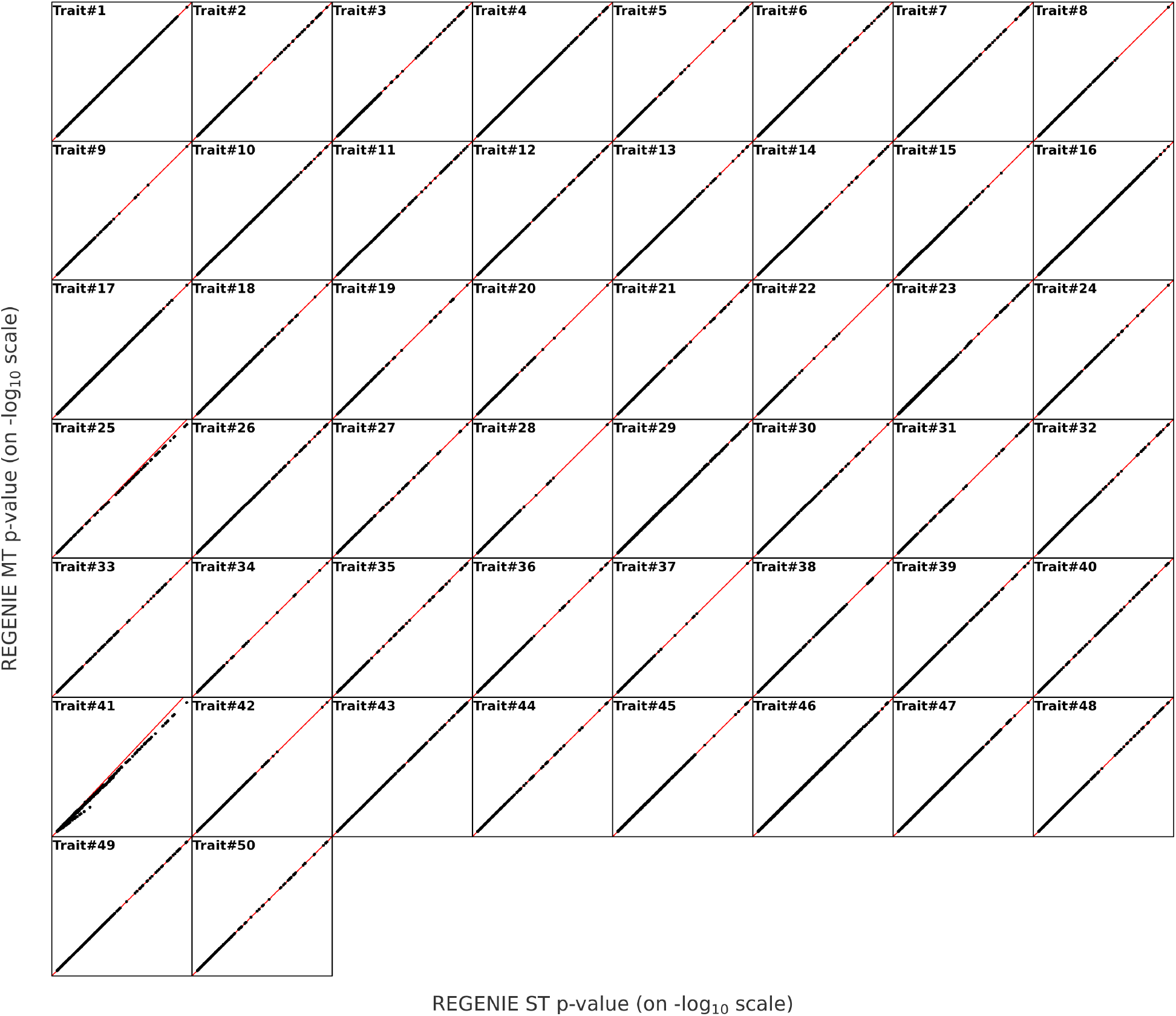
Evaluation of REGENIE’s multi-trait missing data approximation for quantitative traits. Scatter-plots comparing association results with REGENIE for 50 quantitative traits using UK Biobank white British samples. Both step 1 and 2 of REGENIE are ran once for all 50 traits (REGENIE MT), and they are also ran separately for each trait (REGENIE ST), where missing observations are dropped from the analysis. 11.6 million imputed SNPs are tested for association with each trait. Information on the traits is available in **Supplementary Table S2**. The scaling of the axes varies over the 50 traits.

**Figure S18:**
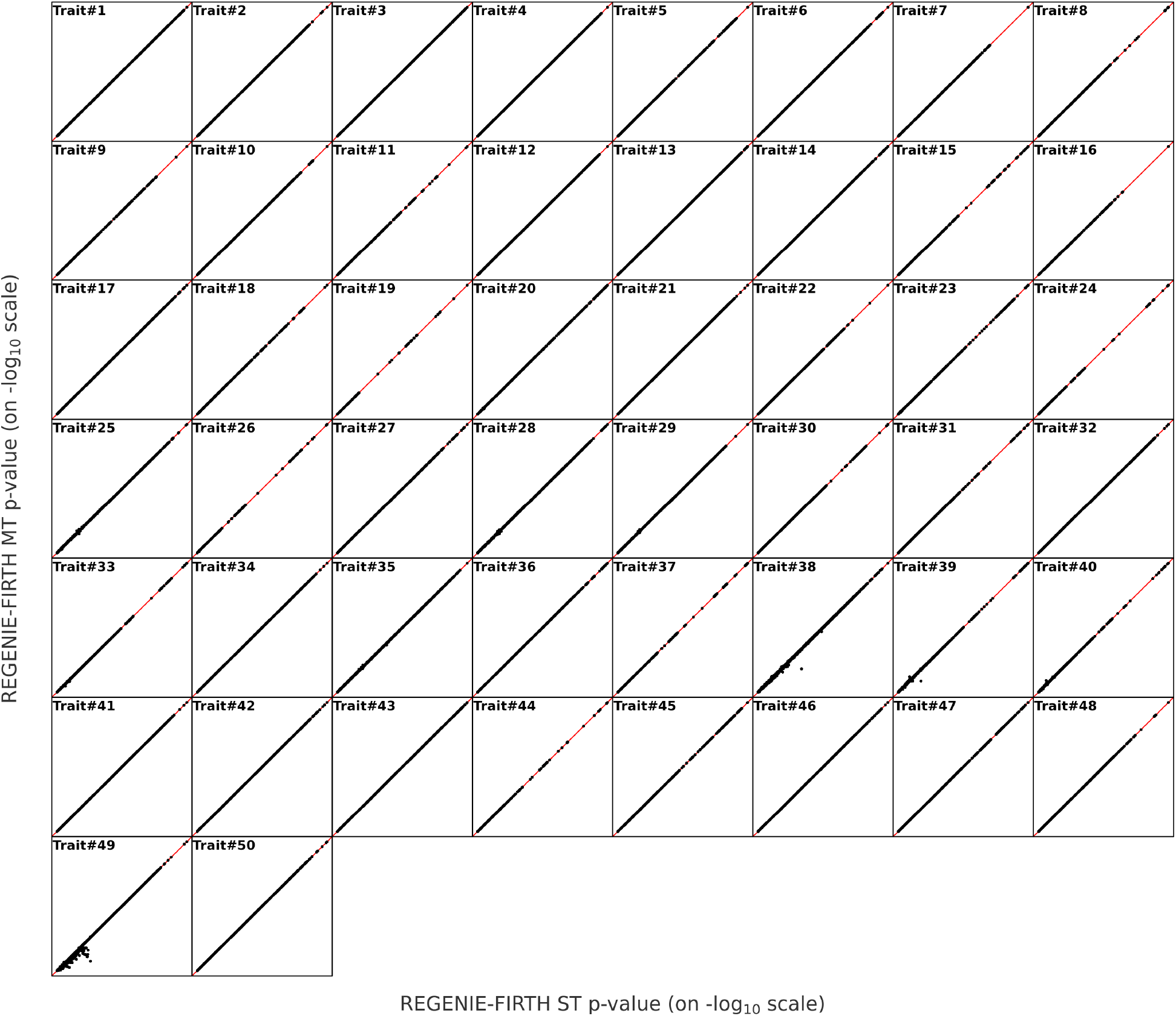
Evaluation of REGENIE’s multi-trait missing data approximation when using a Firth test for binary traits. Scatter-plots comparing association results with REGENIE-SPA for 50 binary traits using UK Biobank white British samples. Both step 1 and 2 of REGENIE were run once for all 50 traits (REGENIE-FIRTH MT) shown on the y-axis, and compared to running each trait separately (REGENIE-FIRTH ST), where missing observations are dropped from the analysis on the x-axis. 11 million imputed SNPs are tested for association with each trait and FIRTH correction is used in REGENIE. Information on the 50 traits is available in **Supplementary Table S6**. The scaling of the axes varies over the 50 traits.

**Figure S19:**
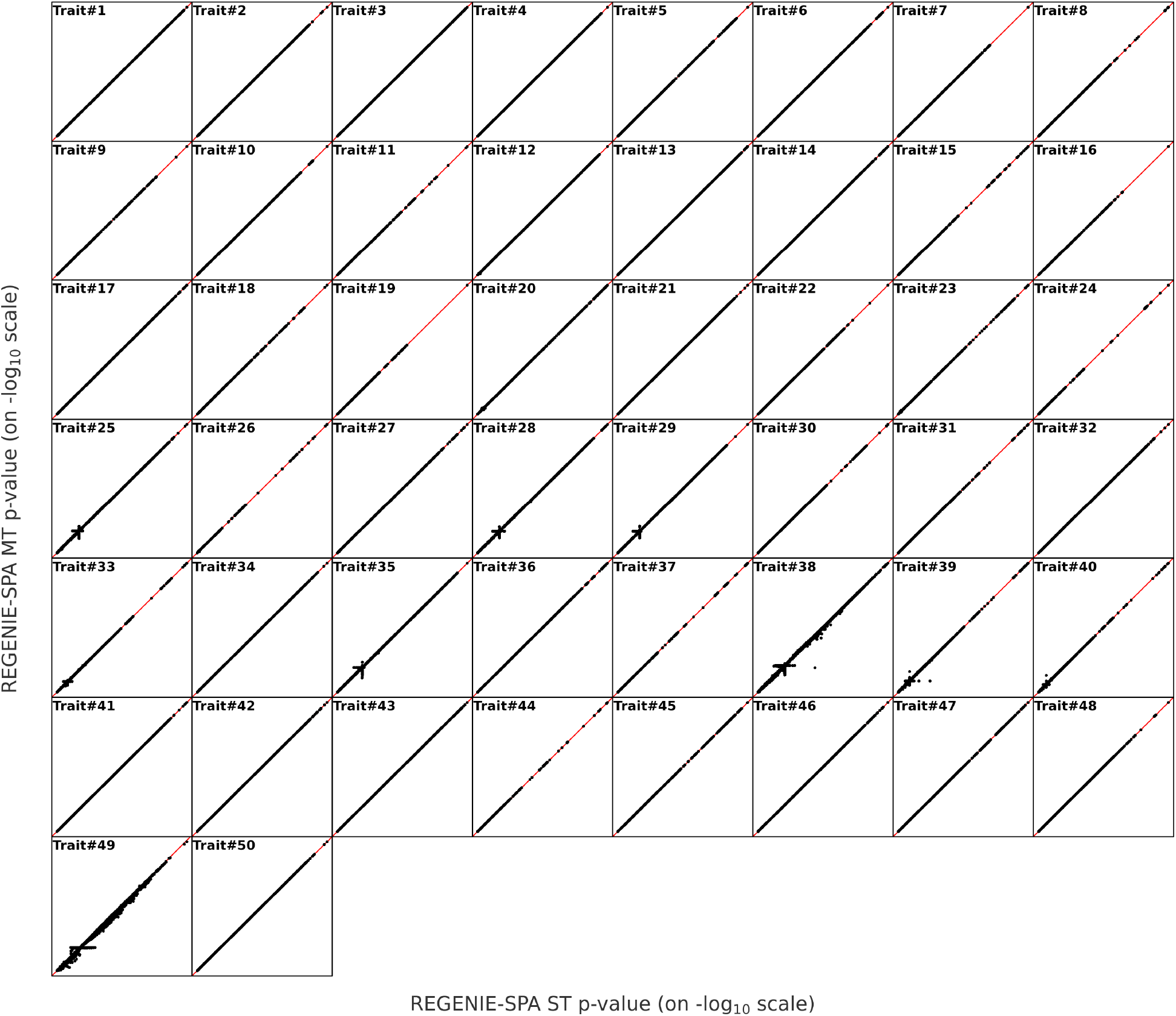
Evaluation of REGENIE’s multi-trait missing data approximation when using a SPA test for binary traits. Scatter-plots comparing association results with REGENIE-SPA for 50 binary traits using UK Biobank white British samples. Both step 1 and 2 of REGENIE were run once for all 50 traits (REGENIE-SPA MT) shown on the y-axis, and compared to running each trait separately (REGENIE-SPA ST), where missing observations are dropped from the analysis on the x-axis. 11 million imputed SNPs are tested for association with each trait and SPA correction is used in REGENIE. Information on the 50 traits is available in **Supplementary Table S6**. The scaling of the axes varies over the 50 traits.

**Figure S20:**
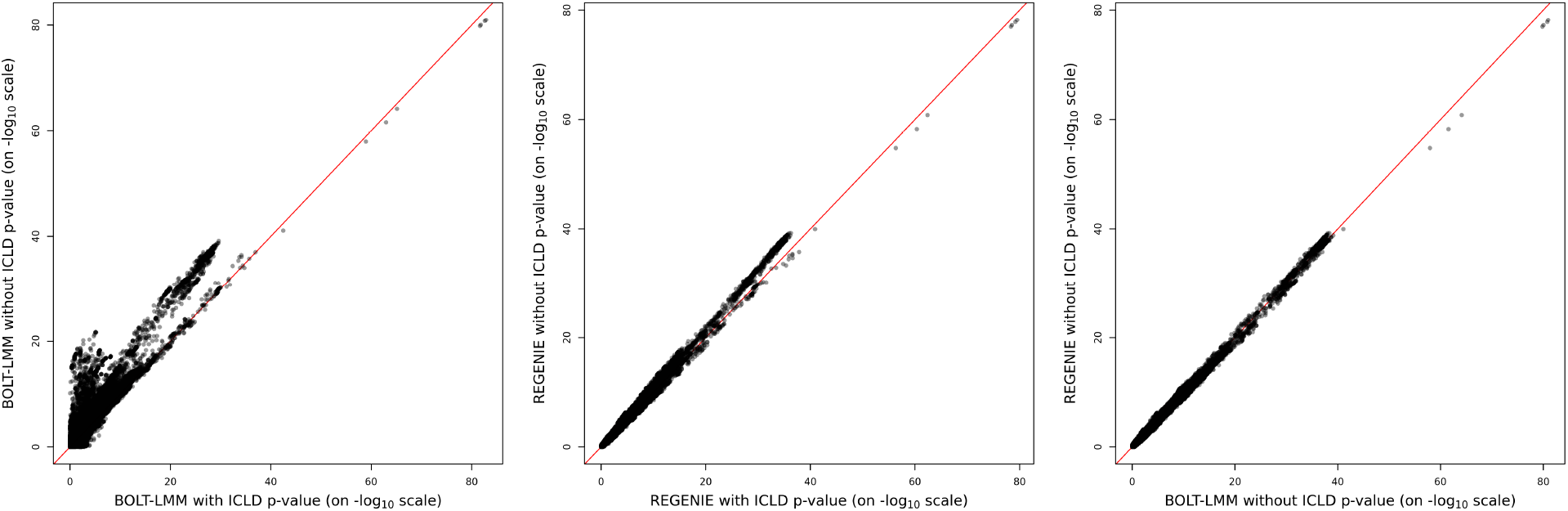
Effect of inter-chromosome LD (ICLD) on different methods. The scatter-plots show association results for LDL in 321,179 UK Biobank white British samples for REGENIE and BOLT-LMM in the presence of inter-chromosome LD (ICLD). Step 1 of REGENIE and BOLT-LMM were run including (with ICLD) and excluding (no ICLD) SNP-pairs involved in inter-chromosomal LD. 745,696 imputed SNPs on chromosome 6 are tested in step 2.

**Figure S21:**
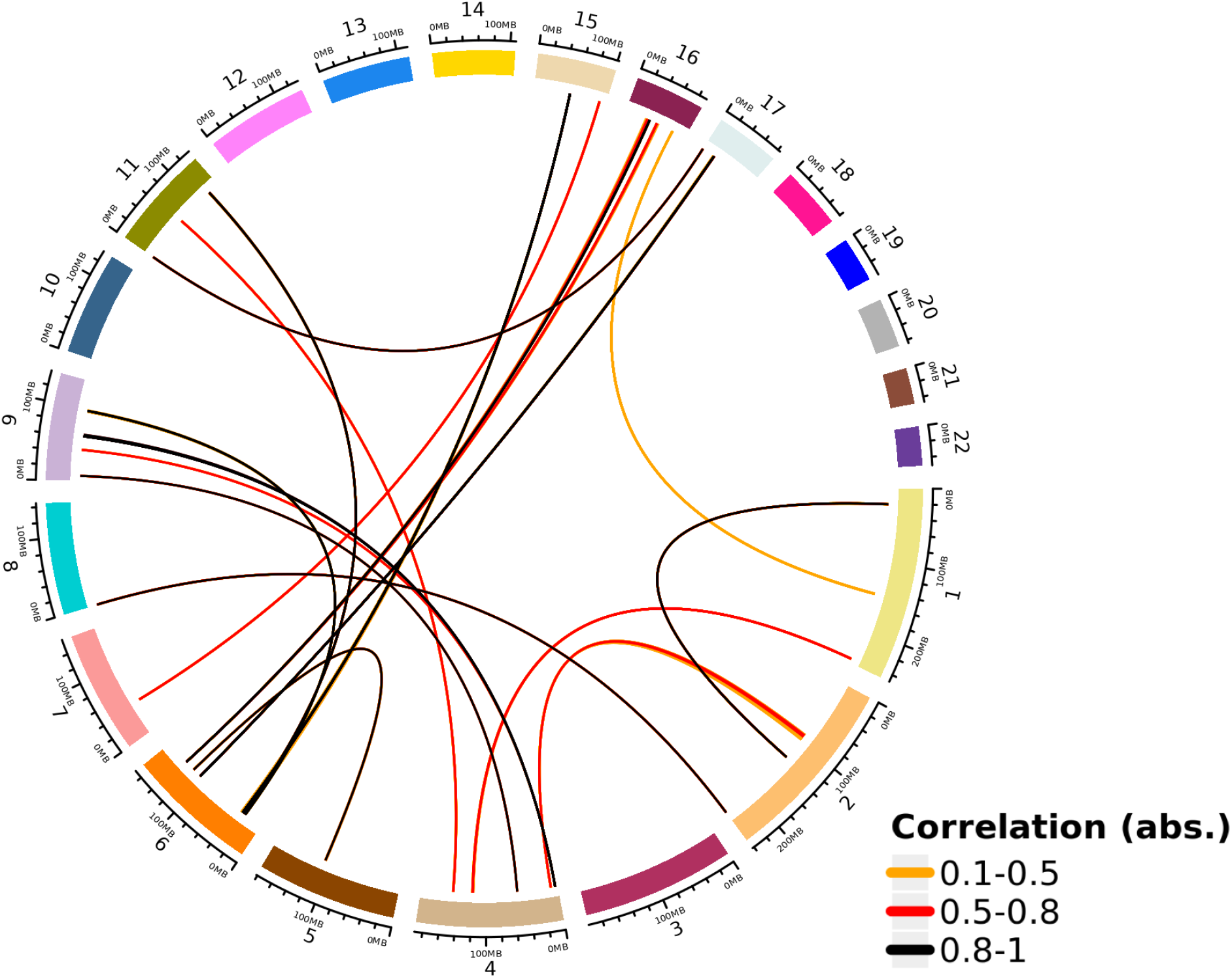
Circle plot of the inter-chromosome LD (ICLD) present in the UK Biobank array data. Each colored region in the perimeter of the circle represents a chromosome. Each link going through the circle indicates the chromosome and physical positions of the pair of SNPs involved in ICLD. The magnitude of the correlation between the SNPs in ICLD is represented by the color of the link. A total of 3,697 SNP-pairs are displayed in the plot (some of the links overlap). The distance between consecutive tick marks represents 20Mb.

